# *Pseudomonas aeruginosa* isolates defective in function of the LasR quorum sensing regulator are frequent in diverse environmental niches

**DOI:** 10.1101/2021.03.25.437011

**Authors:** Marie-Christine Groleau, Hélène Taillefer, Antony T. Vincent, Philippe Constant, Eric Déziel

## Abstract

The saprophyte *Pseudomonas aeruginosa* is a versatile opportunistic pathogen causing infections in immunocompromised individuals. To facilitate its adaptation to a large variety of niches, this bacterium exploits population density-dependant gene regulation systems called quorum sensing. In *P. aeruginosa*, three distinct but interrelated quorum sensing systems (*las*, *rhl* and *pqs*) regulate the production of many survival and virulence functions. In prototypical strains, the *las* system, through its transcriptional regulator LasR, is important for the full activation of the *rhl* and *pqs* systems. Still, LasR-deficient isolates have been reported, mostly sampled from the lungs of people with cystic fibrosis, where they are considered selected by the chronic infection environment. In this study, we show that a defect in LasR activity appears to be an actually widespread mechanism of adaptation in this bacterium. Indeed, we found abundant LasR-defective isolates sampled from hydrocarbon-contaminated soils, hospital sink drains, and meat/fish market environments, using an approach based on phenotypic profiling, supported by gene sequencing. Interestingly, several LasR-defective isolates maintain an active *rhl* system or are deficient in *pqs* system signaling. The high prevalence of a LasR-defective phenotype among environmental *P. aeruginosa* isolates questions the role of quorum sensing in niche adaptation.

## INTRODUCTION

Many Gram-negative bacteria found in natural habitats, such as members of the *Pseudomonas* genus, are also capable of causing opportunistic infections in plants and animals, including humans. Clinical and environmental strains of *Pseudomonas aeruginosa* possess a high degree of genomic conservation but still show significant phenotypic diversity (1). This opportunistic pathogen is an adaptable organism mostly distributed in environments closely related to human activities and is a major cause of nosocomial infections (2). To promote survival in various environmental niches, *P. aeruginosa* coordinates group behaviors to act as a community through density-dependent intercellular communication systems named “quorum sensing” (QS). QS is important for the coordination of competitive and cooperative interactions between organisms within a species or between species (3). In *P. aeruginosa*, the QS circuitry is built on three hierarchically arranged systems, each mediated by the production and detection of distinct signaling molecules. The *las* system is generally considered to initiate the regulatory cascade (4). LasI is a LuxI-type synthase responsible for production of *N*-3-oxododecanyol-homoserine lactone (3-oxo-C_12_-HSL), the signaling molecule that binds and activates the LuxR-type transcriptional regulator LasR. LasR then induces the transcription of many target genes including those encoding various exoproteases (e.g., LasA, LasB) as well as the *lasI* gene, thus causing a positive feedback loop (5–7). LasR bound to its autoinducer 3-oxo-C_12_-HSL also activates the downstream QS system *rhl*, by inducing the transcription of both *rhlI* and *rhlR* (4, 7–10). The RhlI synthase catalyzes the production of *N*-butanoyl-homoserine lactone (C_4_-HSL) that can bind to the LuxR-type regulator RhlR, which controls the transcription of other genes, several encoding for exoproducts such as phenazines (*phz1* and *phz2* operons), rhamnolipids (*rhlAB*) and hydrogen cyanide (*hcnABC*) (8, 11–14).

The third QS system is based on the production of 4-hydroxy-2-alkylquinolines (HAQs), which are synthesized through the enzymes encoded by the *pqsABCDE* operon. The main product of the PqsABCD enzymes, 4-hydroxy-2-heptylquinoline (HHQ) is converted in the *Pseudomonas* quinolone signal (PQS) by the PqsH enzyme (15). Both HHQ and PQS act as autoinducing ligands of the LysR-type transcriptional regulator MvfR (PqsR), which in turn activates the transcription of the *pqs* operon. The PqsABCD enzymes coupled with the activity of the PqsL monooxygenase also synthetize 4-hydroxy-2-heptylquinoline *N*-oxide (HQNO), a secondary metabolite inhibiting Gram-positive bacteria respiration and inducing self-poisoning (16–18). Considering that LasR also controls the transcription of both the *pqsH* and *pqsL* genes (7), the *pqs* system is in fact intertwined in the QS circuitry, with LasR positively regulating the expression of *pqsABCDE* and RhlR acting as a repressor. Furthermore, the PqsE protein is not required for HAQ synthesis but is important for full activation of RhlR-dependent QS (19, 20). Importantly, the above-mentioned hierarchy of the QS circuitry is flexible, since RhlR can take over the role of LasR in the stationary phase (21), and several clinically isolated LasR-defective mutants are still able to produce RhlR-dependent factors (22, 23).

LasR-defective mutants are frequently identified among clinical samples, especially from chronic infections such as cystic fibrosis (23–27). The presence of these mutants is associated with an adaptation to unique conditions. For example, they possess growth advantages on specific amino acids largely found in CF secretions or have an increased resistance to some antimicrobials, suggesting a selective pressure from this kind of environment (25, 26). On the other hand, LasR-defective mutants were also exceptionally isolated from corneal ulcers, considered an acute infection (28). Surprisingly, the conservation of QS activity has been rarely investigated among isolates from environmental sources (27, 29). LasR-deficient isolates of *P. aeruginosa* have been thoroughly characterized in several reports, but essentially in studies investigating their prevalence in the lungs of patients with cystic fibrosis (CF) (23, 26). Focus is often on the findings of variations in the sequence of the *lasR* gene. However, sequences are compared to those of reference strains and variations may not always indicate a defective LasR activity. Also, variations in regulatory elements affecting the functionality of LasR are set aside in these studies, even more so when focusing on the sole sequencing of the *lasR* gene rather than whole genome sequencing. Elements such as anti-activator proteins such as QslA, QteE and QscR are able to suppress QS-activation trough the blocking of LasR activity (30–33). A phage-encoded anti-activator named Aqs1 that inhibits LasR activity was also described (34). Phenotypical traits of LasR-defective strains are well known. A non-functional LasR typically leads to a characteristic surface iridescent sheen and autolysis of colonies on agar plates, a result of imbalance in HAQ production (26). The production of the blue pigment pyocyanin is also negatively altered in exponential phase cultures grown in rich media. On the other hand, pyocyanin accumulates at high levels in cultures of LasR-deficient strains during the late stationary phase (21, 35, 36). Since LasR controls the production of various proteases, a LasR deficiency also leads to an inability to degrade casein (5, 6).

In this study, we tested the hypothesis that a defect in LasR activity is not restricted to clinical strains originating from human infections. To estimate the prevalence of a LasR defect in a collection of strains isolated from various environments other than clinical, we chose an approach based on phenotypic profiling rather than strictly on gene sequencing. The phenotypic profile of 176 environmental isolates originating from hospital sink drains, hydrocarbon-contaminated soils and meat/fish market environment and products was characterized and integrated into a classification scheme providing non-ambiguous assessment of their LasR activity status. The classification approach was further supported by sequencing of selected isolates. Unexpectedly, we found that LasR-defective isolates are ubiquitous in these environments. We also found an abundance of LasR-defective isolates maintaining an active *rhl* system as well as isolates with a functional LasR but producing no HAQ molecules. This study thus reveals a high prevalence of the LasR-defective phenotype among *P. aeruginosa* natural isolates. Their incidence in CF infections is associated with their selection by the chronic infection environment. Rather, loss of LasR activity appears a widespread mechanism of adaptation in this bacterium.

## MATERIALS AND METHODS

### Strains and growth conditions

The 174 isolates included in the analysis were collected in Canada and Ivory Coast. In Canada, samples were collected from an oil-contaminated sand pit (37) and from the environment (tap aerator, drain, sink surface) of 18 sinks from nine different rooms in an hospital (38), both in the Montreal region. In Ivory Coast, samples were collected from bovine meats, fresh or smoked fish, and their direct environment (stalls, cutting boards) in five different public markets in Abidjan (39). We also included isolates PUPa3 and PG201 isolated from soils in India and Switzerland, respectively (40, 41). A detailed list of the isolates and their origin is presented in **Table S1**. Bacteria were routinely grown in Lysogeny broth (LB) (AlphaBiosciences, USA) or Tryptic Soy Broth (TSB, Difco) at 37°C in a TC-7 Roller Drum (New Brunswick) at 150 rpm. Reference strains were PA14 (42), its isogenic *lasR*::Gm mutant (43) and clinical *lasR* mutant isolate E90 (23).

### Phenotypical assays

For pyocyanin measurements, bacteria were grown in King’s A broth with 10 μM FeCl_3_ (44) for 18h at 37°C. Briefly, an overnight culture in TSB was diluted to an OD_600_ of 0.01. At chosen times during growth, a sample of the culture was centrifuged (10,000 x *g*, 15 min). The supernatant was transferred to a 96-well plate and the OD_695_ was measured using a Cytation microplate reader (BioTek, VT, USA). Since clumping of cells was apparent in cultures of many strains, we assessed growth using total protein quantification of the cell pellet harvested from the whole culture at the time of sampling. The cell pellets were suspended in 0.1 N NaOH and heated at 70°C for 1 h with frequent vortexing. Protein concentrations were determined using the Bradford quantification assay (Bio-Rad, Canada) with Bovine Serum Albumin as a standard.

To detect iridescence and autolysis of colonies, bacteria were grown on LB agar plates for 24h at 37°C and colonies were visually inspected for the appearance of a metallic sheen (iridescence) and the presence of autolysis zones.

Protease production was ascribed to strains displaying the ability to grow on casein as a sole carbon source. Bacteria were grown overnight in LB broth. Five microliters of culture were deposited on the surface of M9 media agar containing 0.1% sodium caseinate as sole carbon source. The plates were incubated at 37°C for 24-48h before visual inspection.

### Quantification of quorum sensing signaling molecules

Concentrations of HAQs and AHLs in cultures were measured for bacteria grown in King’s A medium. Based on their production profile, concentrations of 3-oxo-C_12_-HSL, C_4_-HSL were measured at 3h (1) and 6h (2) time points. The main C_7_ HAQ congeners HHQ, HQNO and PQS were measured at 6h (1) and 24h (2). Briefly, an equal volume of methanol containing internal standard tetradeuterated 4-hydroxy-2-heptylquinoline (HHQ-d4) was added to the culture sample. The suspension was vortexed and centrifuged 5 min at maximum speed to remove bacteria. The resulting supernatant was transferred in vials for liquid chromatography/mass spectrometry/mass spectrometry (LC/MS/MS) analyses, as previously reported (45) For quantifications of AHLs, ethyl acetate extracts were analyzed by LC/MS/MS as described (45). The cell pellets were suspended in 0.1N NaOH and heated at 70°C for 1 hour with frequent vortexing. Protein concentrations were obtained using the Bradford quantification assay (BioRad). Production (mg/L) is presented as a ratios over biomass (μg/mL of proteins).

### Measurement of the expression of *rhlA-gfp*

Bacteria were transformed by electroporation with plasmid pJF01 containing the promoter region of the *rhlA* gene in front of the *gfp* reporter gene (23). Transformants were selected on 30 μg/mL gentamicin. To evaluate the expression of the *rhlA*-*gfp* reporter, bacteria were grown overnight in TSB containing 30 μg/mL gentamicin. After three washes with PBS, cells were suspended in King’s A media and transferred in a 96-well microplate. The plate was incubated for 30 hours at 37°C with shaking. Fluorescence was measured (excitation: 489 nm, emission: 520 nm) using a Cytation microplate reader (BioTek, VT, USA). Relative fluorescence units (RFU) data were normalized by the OD_600_.

### Sequence analyses

The genomic sequences of 45 of our environmental *P. aeruginosa* strains available in GenBank (**Table S4**) were compared to the reference strain UCBPP-PA14 (GenBank: CP000438.1) using snippy version 4.6.0 (https://github.com/tseemann/snippy).

### Statistical analyses

Statistical analyses were performed using the software R (46). Pairwise comparison of phenotypic profiles was expressed under the Euclidean distance units computed on Hellinger-transformed molecule concentrations matrix. Distance matrix was represented with dendrogram elaborated with agglomerative hierarchical clustering based on the Ward’s linkage method. Cluster stability was assessed by bootstrap resampling method (47) implemented in the package ‘fcp’. The heatmap, the dendrogram were done with the package ‘ggplot2’(48). The most distinctive variables amongst stains were identified with a principal component analysis (PCA). All raw data used for clustering analyses are found in **Table S3**.

## RESULTS

### Many environmental *Pseudomonas aeruginosa* isolates are deficient in LasR activity

To investigate the prevalence of a LasR deficiency among non-clinical *P. aeruginosa* isolates, and since sequencing is not sufficient to identify a defect in LasR activity, we performed a phenotypic characterization of 176 isolates sampled from diverse environments (**Table S1**), based on known traits of LasR-defective strains. We first looked at unique phenotypes (iridescence and autolysis) of colony on agar, at pyocyanin production patterns in both liquid TSB and King’s A medium, and at the ability to grow on casein as a sole carbon source, indicative of exoprotease production. As controls, PA14 strain and its isogenic *lasR*-null mutant were used. Results are detailed in **Table S2**. Based on these often used criteria, we were unable to clearly determine which isolates had a LasR deficiency. We observed variations in the ability to grow on casein; inconsistencies in the correlation between factors, for instance some isolates were iridescent and autolytic, overproduced pyocyanin in King’s A but could still grow on casein agar. It appears that these traits, although typically used as markers of LasR functionality, are not reliable enough for efficient determination of LasR deficiency in phenotypically-diverse environmental isolates, since there is no specific threshold delineating this characteristic.

We thus looked for an alternative approach. LasR controls the production of extracellular metabolites such as AHLs, HAQs and pyocyanin, whose concentration profiles over time could be adapted to a classification scheme to establish a LasR phenotype. That strategy was explored by measuring 3-oxo-C_12_-HSL, C_4_-HSL as well as HHQ, HQNO, PQS and pyocyanin concentrations at various time points during growth of a *lasR* mutant of PA14, to define reference production profiles. Biosynthesis of these metabolites is directly or indirectly dependent on the regulatory activity of LasR. In the *lasR* mutant background, the production of C_4_-HSL was not observed until early stationary phase due to the delayed activation of *rhlI* by RhlR whereas there is absence of 3-oxo-C_12_-HSL (**Figs. 1a and b**) (20). Also, HQNO and PQS are expectedly only produced in minute levels in a LasR-deficient mutant (**Figs. 1e and f**). Accordingly, the mutant accumulates HHQ, the precursor of PQS (**Fig. 1d**) since in wild-type PA14, the levels of HHQ stabilize when concentrations of HQNO and PQS start augmenting (**Figs. 1d, e and f**) (15). As expected, pyocyanin production is heightened in the *lasR*::Gm mutant when compared to PA14 (**Fig. 1c**). Taken together, the distinct production profiles of all these metabolites suggest that they could be used as markers of LasR deficiency.

**Figure 1.**
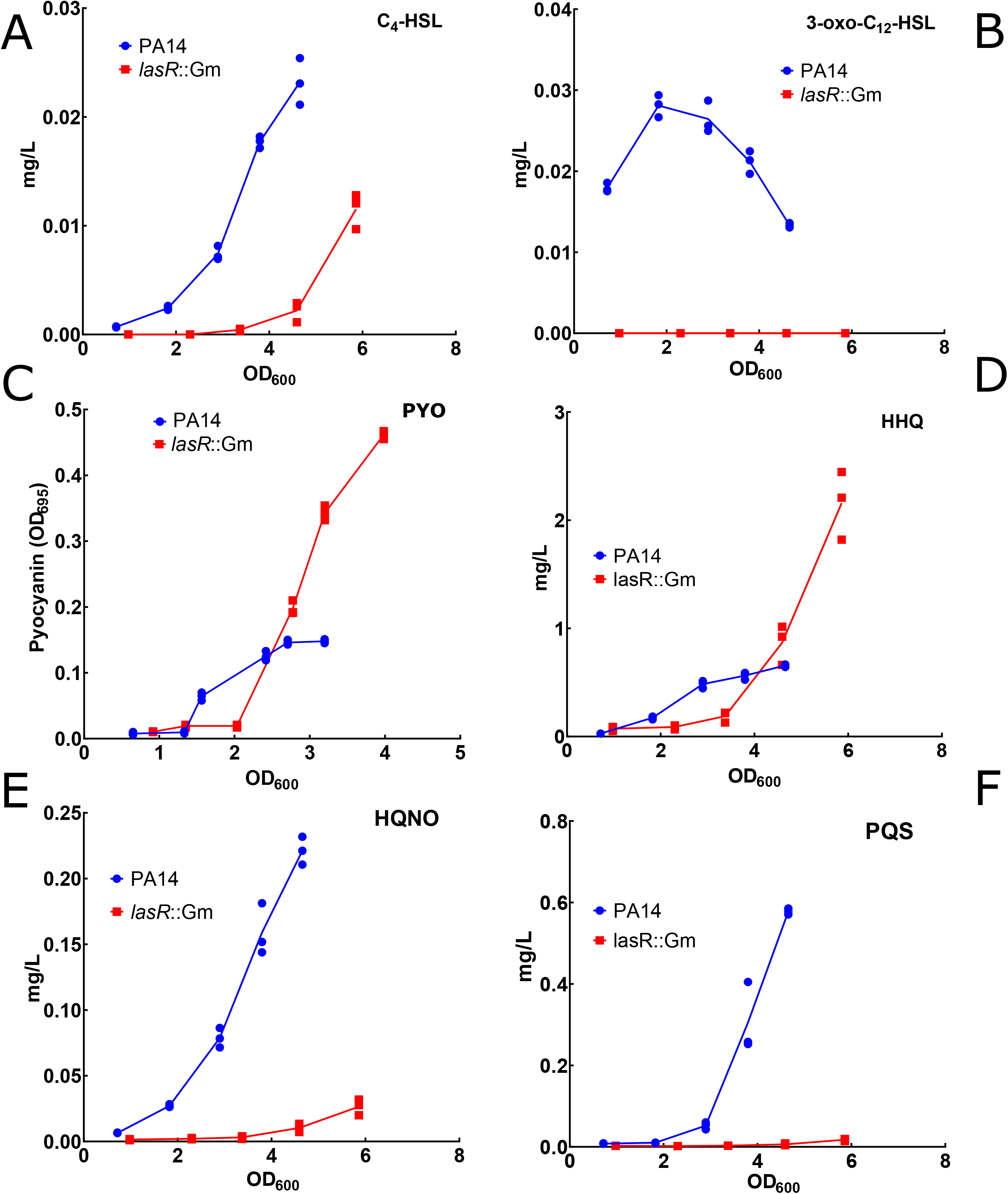
Production profiles of various extracellular metabolites in strain PA14 and its isogenic *lasR*-mutant. 3-oxo-C_12_-HSL, C_4_-HSL, HHQ, PQS, HQNO were measured by LC/MS and pyocyanin (PYO) production assessed using A_695_ of the culture supernatant at various time points during growth. Cultures were prepared in triplicates in TSB and incubated à 37°C.

Convergence of metabolic profiles with the LasR-defective phenotype was further examined with a subset of 30 isolates from various environments. Based on our initial qualitative phenotypic profiling (**Table S2**), we purposely chose a majority of isolates with profiles possibly corresponding to a LasR-deficiency as well as some with patterns in accordance with a functional LasR. PA14 and its *lasR*- mutant were included as controls. Timing of sampling was chosen based on the profiles observed in the LasR mutant vs WT PA14 (**Fig. 1**). AHLs were sampled after 3h and 6h which correspond to OD_600_ of 1 and 3, respectively, to generate variables C4-HSL_1 and _2 and 3-oxo-C12-HSL_1 and _2. HHQ, HQNO and PQS were sampled at 6h and 24h to generate variables HHQ_1 and _2, HQNO_1 and _2, PQS_1 and _2, respectively. Pyocyanin was measured at 24h (PYOKA_2). A Principal Component Analysis (PCA) was performed to examine variables contributing the most in distinguishing examined strains and identify redundancy consistency between certain traits (**Fig. 2**). This led to the elaboration of a parsimonious classification scheme integrating levels of HHQ, HQNO, PQS and pyocyanin in culture samples to discriminate between strains with a deficient or functional LasR regulator. For instance, we found that the variable HHQ.C7_2, which corresponds to concentrations of HHQ at late stationary phase (24h time point), can explain the distribution of strains as well as AHL concentration data, since they show inverse directions on the same axis (**Fig. 2**). It is noteworthy that there are methodological benefits in avoiding quantifying AHLs from cultures on a large collection of isolates, as these molecules as present in lower concentrations and can be unstable. Moreover, classification of our selected isolates performed using only both AHLs (C4-HSL_1 and _2 and 3-oxo-C12-HSL_1 and _2) displayed the same pattern as with only HAQs and pyocyanin, confirming that our choice of variables is suitable (data not shown).

**Figure 2.**
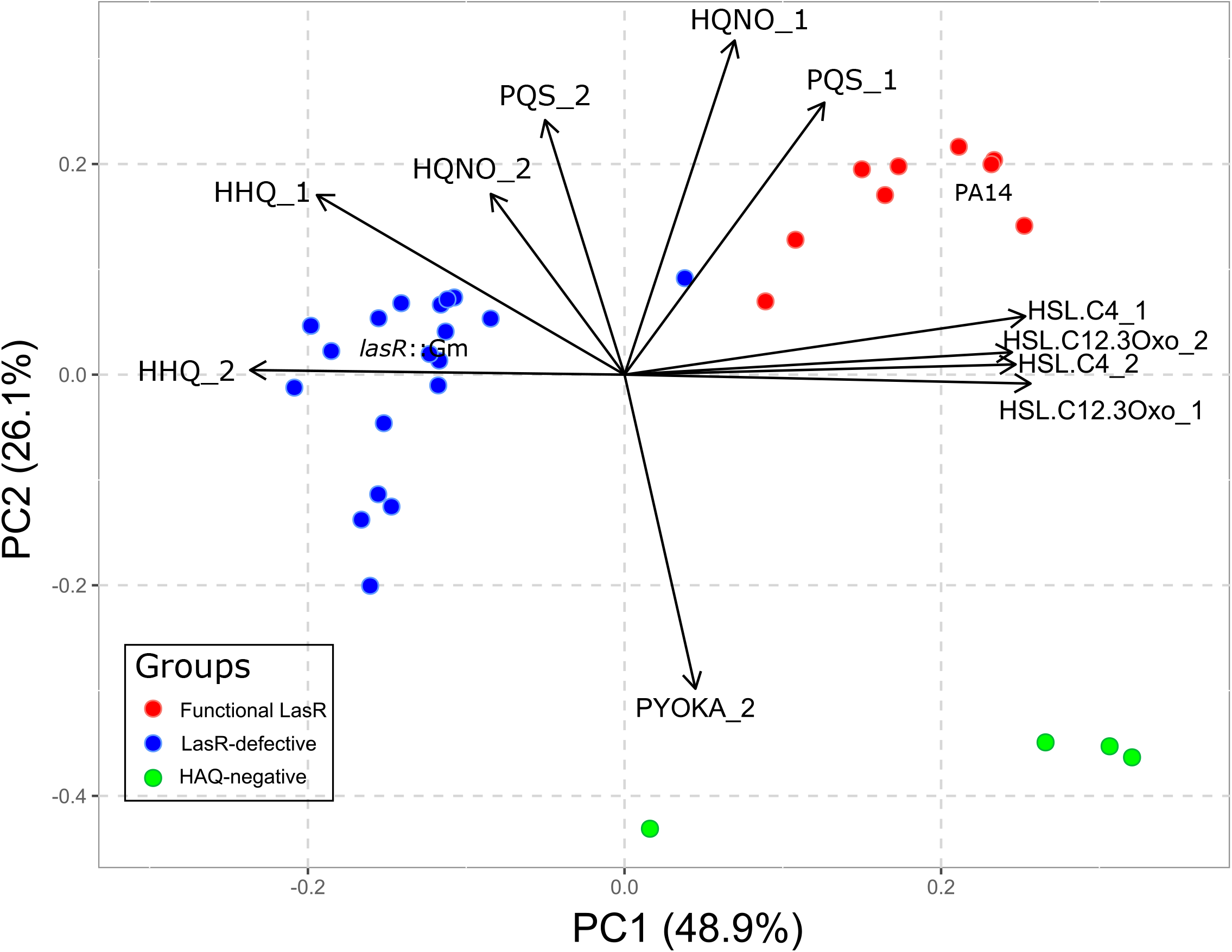
Principal component analysis elaborated with levels of extracellular metabolites analyzed from cultures of a subset of *Pseudomonas aeruginosa* environmental isolates sampled at two time points. Concentrations of C_4_-HSL and 3-oxo-C_12_-HSL were measured at 3h (C4-HSL_1 and 3-oxo-C12-HSL_1) and 6h (C4-HSL_2 and 3-oxo-C12-HSL_2), respectively. Concentrations of HHQ, HQNO, and PQS sampled at 6h (HHQ_1, HQNO_1, PQS_1) and 24h (HHQ_2, HQNO_2, PQS_2), respectively. Pyocyanin (PYOKA_2) was sampled at 24h. Strain PA14 and its isogenic *lasR* mutant are included as controls. Groups represent clusters based on a dendrogram elaborated with the same variables (**Fig. S1**).

According to pairwise comparison of the strains based on our parsimonious model, three different phenotypic profiles were identified among our subset of 30 isolates (**Fig. 3**). The functional LasR group, behaving typically like WT PA14, encompasses isolates which produce both PQS and HQNO at the end of the exponential phase (PQS.C7_1 and HQNO.C7_1). The LasR-defective group is represented by isolates displaying high concentrations of HHQ at both time points coupled with a delayed production of PQS and HQNO and higher late production of pyocyanin in King’s A, as illustrated in **Figure 1**. Our analysis integrating HAQs and AHLs signals revealed a new, unexpected third group of QS variants which produced very low levels of HHQ, PQS and HQNO but were still able to produce pyocyanin in King’s A medium (PA-CL504, 32SB, DCB22 and PA-CL507). Three of these isolates (PA-CL504, PA-CL507 and DCB22) produce AHLs like a WT, meaning their LasR is functional and instead pointing to a defect in the regulation or biosynthesis of HAQs. The other isolate (32SB) produces no 3-oxo-C_12_-HSL or C_4_-HSL at both early and late exponential phases, suggesting it is LasR-defective (**Table S3**). This explains the ambiguous data of **Table S2**, where this strain has features of a *lasR* mutant but lack iridescence and autolysis, features which depend on HAQ production.

**Figure 3.**
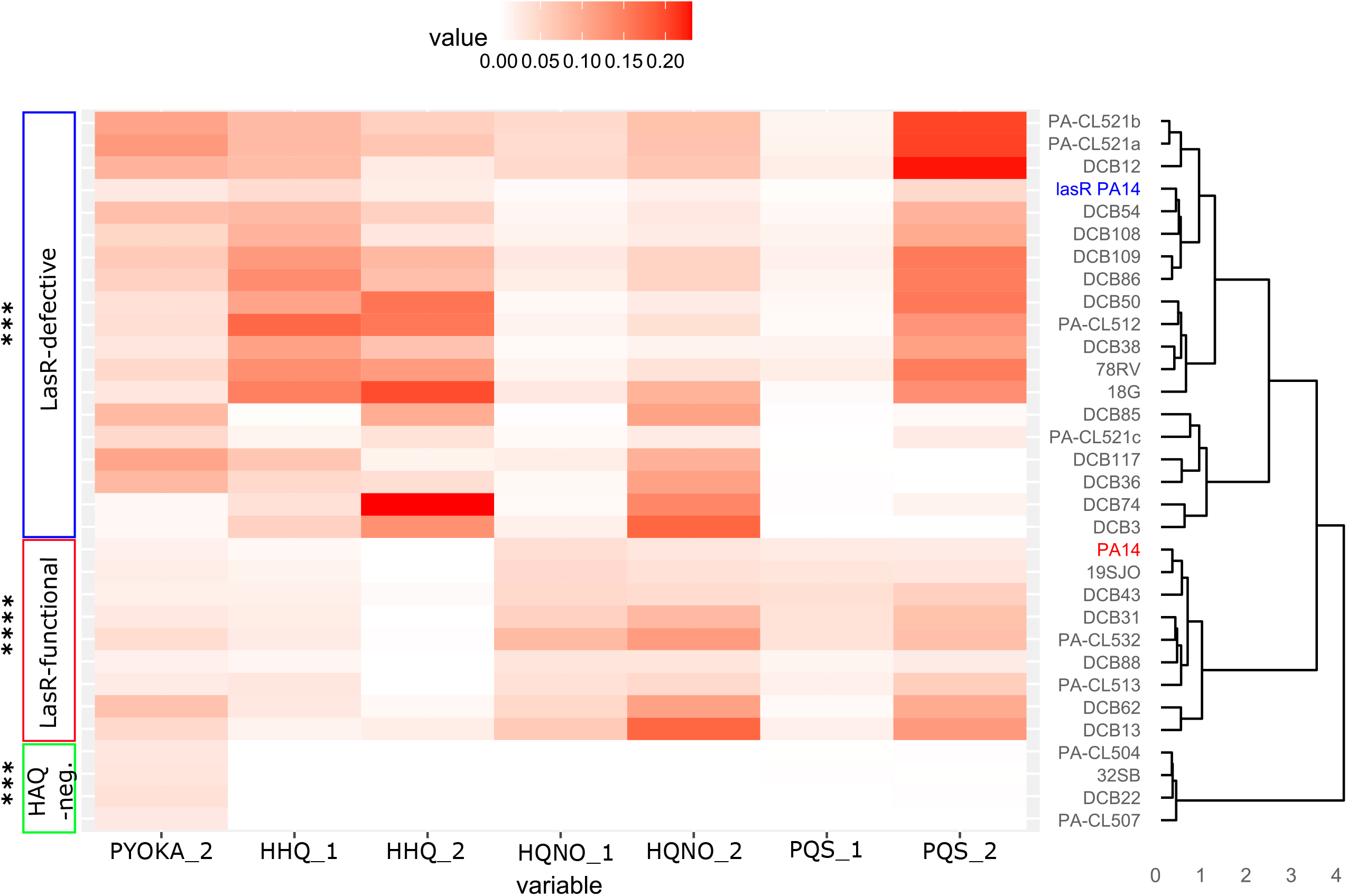
Clustering of LasR-defective isolates. A cluster dendrogram with a heat map was performed on a subset of isolates using chosen variables HHQ_1 and 2, HQNO_1 and 2, PQS_1 and 2 and pyocyanin (PYOKA_1). Concentrations of HHQ_1 and 2, HQNO_1 and 2, PQS_1 and 2 sampled at 6 and 24h. Pyocyanin (PYOKA_2) was sampled at 24h. Strain PA14 and isogenic *lasR* mutant are included as controls. Robustness: **** >90 %, *** 80-89%.

Since our parsimonious classification model including HAQs and pyocyanin production in King’s A allowed us to cluster with good confidence LasR-deficient isolates, we then measured the selected variables for the remaining 146 isolates of our collection. As expected, the strains clustered in three distinct phenotypic groups (**Fig. S2**). One cluster includes PA14-*lasR*::Gm and encompasses isolates with a probable defect in LasR function. The second cluster includes WT PA14 and isolates with a functional LasR. Based on this, at least 40% of our 176 environmental strains display a defective LasR regulatory activity. The third cluster, encompassing 21% of the strains in this collection, contains isolates which produce negligible levels of HAQs. These isolates could also have a defect in their LasR function, thus further increasing the proportion of LasR-defective strains, and/or only be deficient in their ability to synthesize HAQs.

Whole genome sequences are available for 45 of the environmental strains investigated here (**Table S4**) (49). We looked for the presence of mutations in the genomes of these strains compared to PA14 to try to find a genetic explanation for the observed phenotypes. When sequenced isolates had LasR-defective profiles, we indeed found variations in the *lasR* gene when comparing to the *lasR* gene of WT PA14, with a few exceptions (**Table 1**). Validating our initial assumption that sequencing alone cannot predict the LasR phenotype, 19SJV has a typical LasR-defective profile but no mutation in the *lasR* gene was found. Inversely, a variation in the *lasR* gene of 34JS was found but this strain behaves as if its LasR is functional. All together, this phenotypic profiling allowed us to highlight for the first time a strong presence of *P. aeruginosa* isolates with defective LasR-dependant signalling in environments not related to human infections.

**Table 1.**
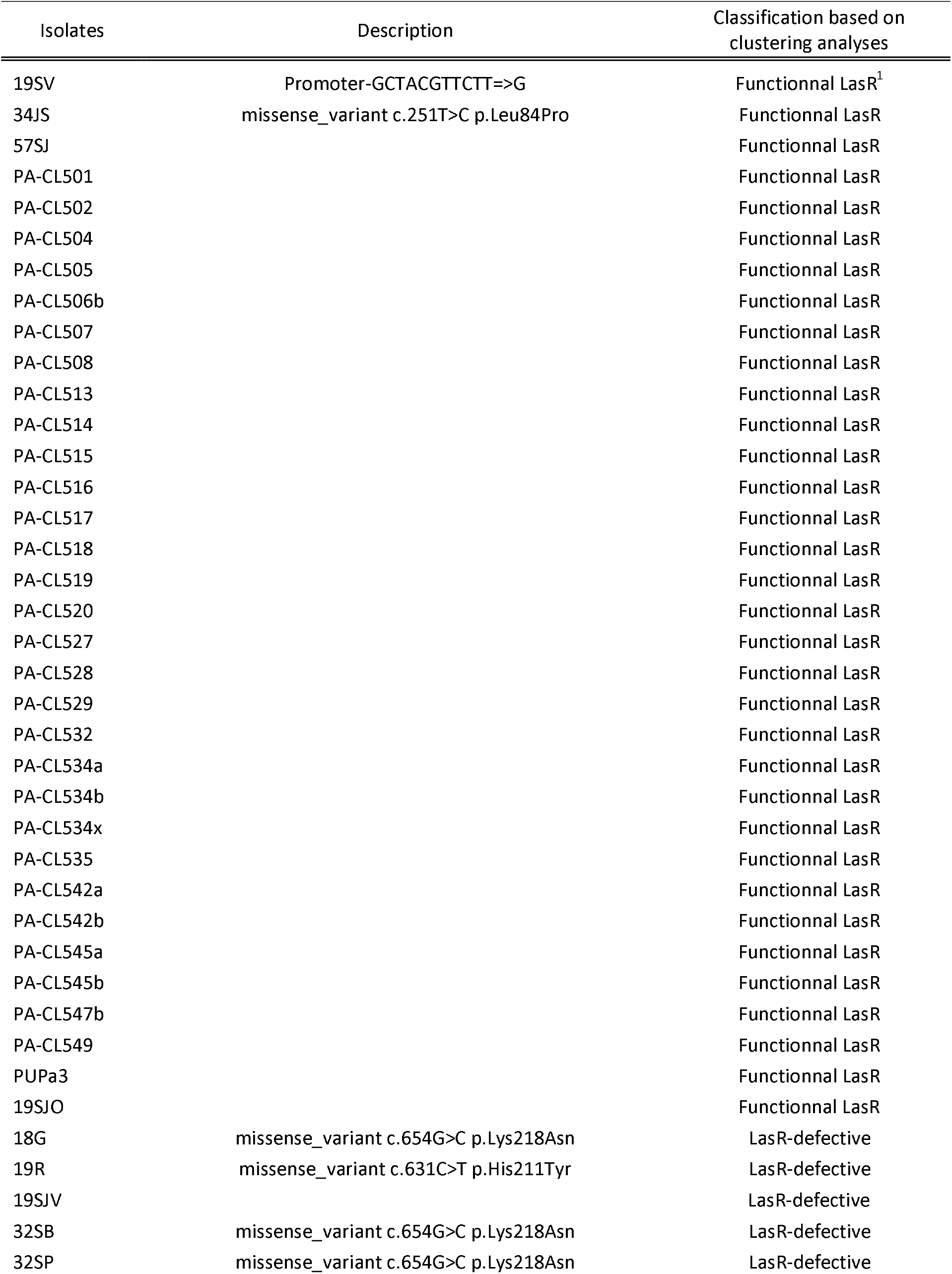
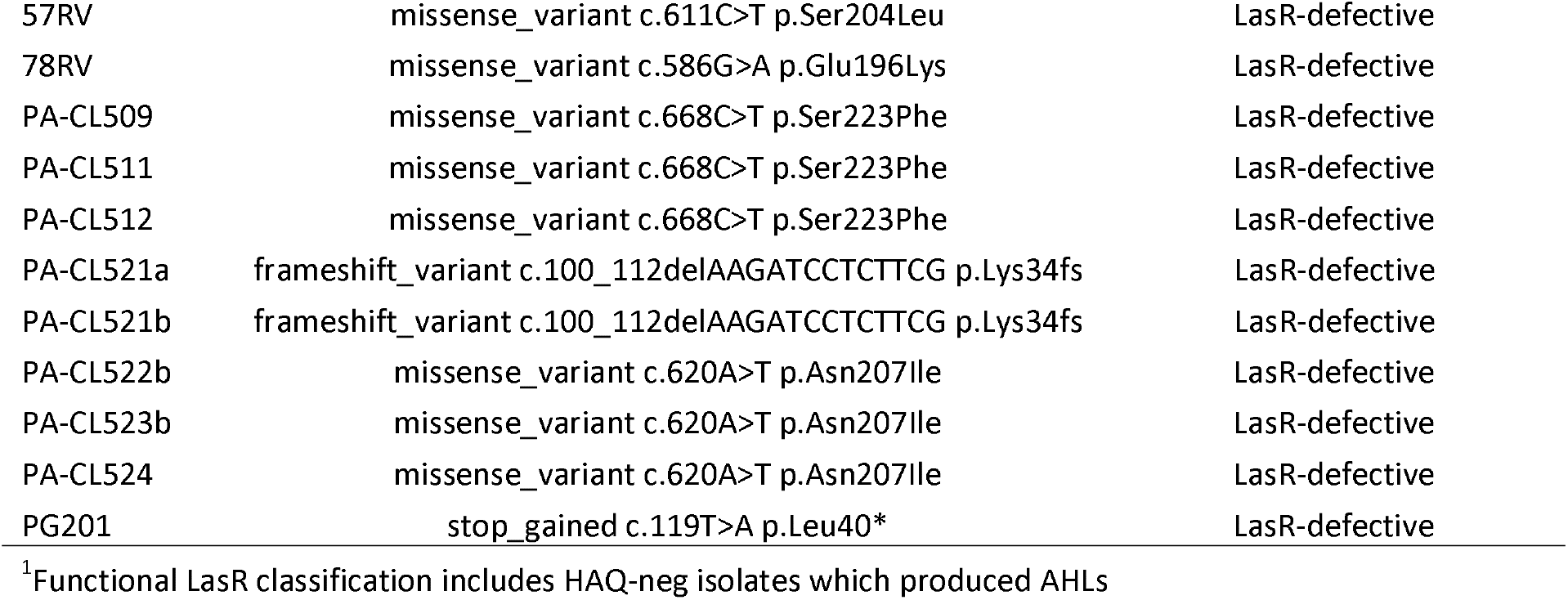
SNPs found in the *lasR* gene of sequenced isolates.

### A subgroup of isolates unable to produce HAQs

As presented above, the third cluster we observed in **Figure 3** contains 38 isolates which produce null or extremely low levels of HAQs. To assess the possibility that some of these isolates could also be LasR-defective, we characterized them further. We measured AHL concentrations and found that 13 of these strains indeed produce both 3-oxo-C_12_-HSL and C_4_-HSL, which indicates a functional LasR (**Table S3**). When these isolates are included in the final evaluation, we conclude that 48% of our 176 isolates are LasR-defective (**Table S3**).

Surprisingly, some HAQ-negative LasR-positive strains still produce pyocyanin in King’s A (**Tables S4 and S5**). When looking at the available sequences for some of these strains, we found no mutations in *mvfR* or in the *pqsBCDE* genes, which could explain the absence of HAQs. The mechanism explaining the absence of HAQs in these strains is yet to be resolved.

### A subgroup of LasR-deficient isolates still capable of RhlR-dependant quorum sensing

A few recent studies reported that a subset of LasR-deficient CF strains conserve a functional RhlR-regulated QS, without the typical requirement for LasR activation, which we call here RAIL strains (for RhlR Active Independently of LasR) (22, 23, 50). We thus asked whether this type of strains in which AHL-dependant QS does not rely on a functional LasR could be also present among our environmental LasR-defective isolates. We looked at RhlR-dependant variables among the subgroup of isolates belonging to the LasR-defective group (**Fig. 3**). Since our clustering key based on HHQ, PQS, HQNO and PYO was not developed to reveal these special LasR-defective strains, we needed to include additional quantitative phenotypes that would be features of RAIL strains. First, since RhlR is a direct activator of the *rhlAB* operon responsible for production of rhamnolipids in *P. aeruginosa*, we chose to measure the activity of a *rhlA-gfp* reporter in these isolates, since it was used to discriminate RAIL isolates before (23). RhlR also regulates the *phzA1* operon which is responsible for the production of phenazines such as pyocyanin. As a second variable, we also measured pyocyanin levels at early stationary phase based on the hypothesis that isolates able to by-pass the traditional LasR-dependant route for the activation of the *rhl* system and thus phenazine production, would be able to produce pyocyanin earlier than typical LasR-deficient strains. We used the previously well-characterized RAIL strain E90 as a reference (50), and performed a clustering analysis. We found that more than half of the LasR-deficient isolates (those studied in **Figs. 2 and 3**) have probable LasR-independent RhlR activity (**Fig. 4 and Table S3**). As previously reported, isolates in this cluster indeed show higher activation at 30h growth in King’s A medium of the *rhlA-gfp* reporter compared to their LasR-defective counterparts (**Table S3**) (23, 50). The PCA analysis confirmed that this variable was the most determinant for classification of the strains but that the earlier timepoint for pyocyanin was not impactful (**Fig. 4**).

**Figure 4.**
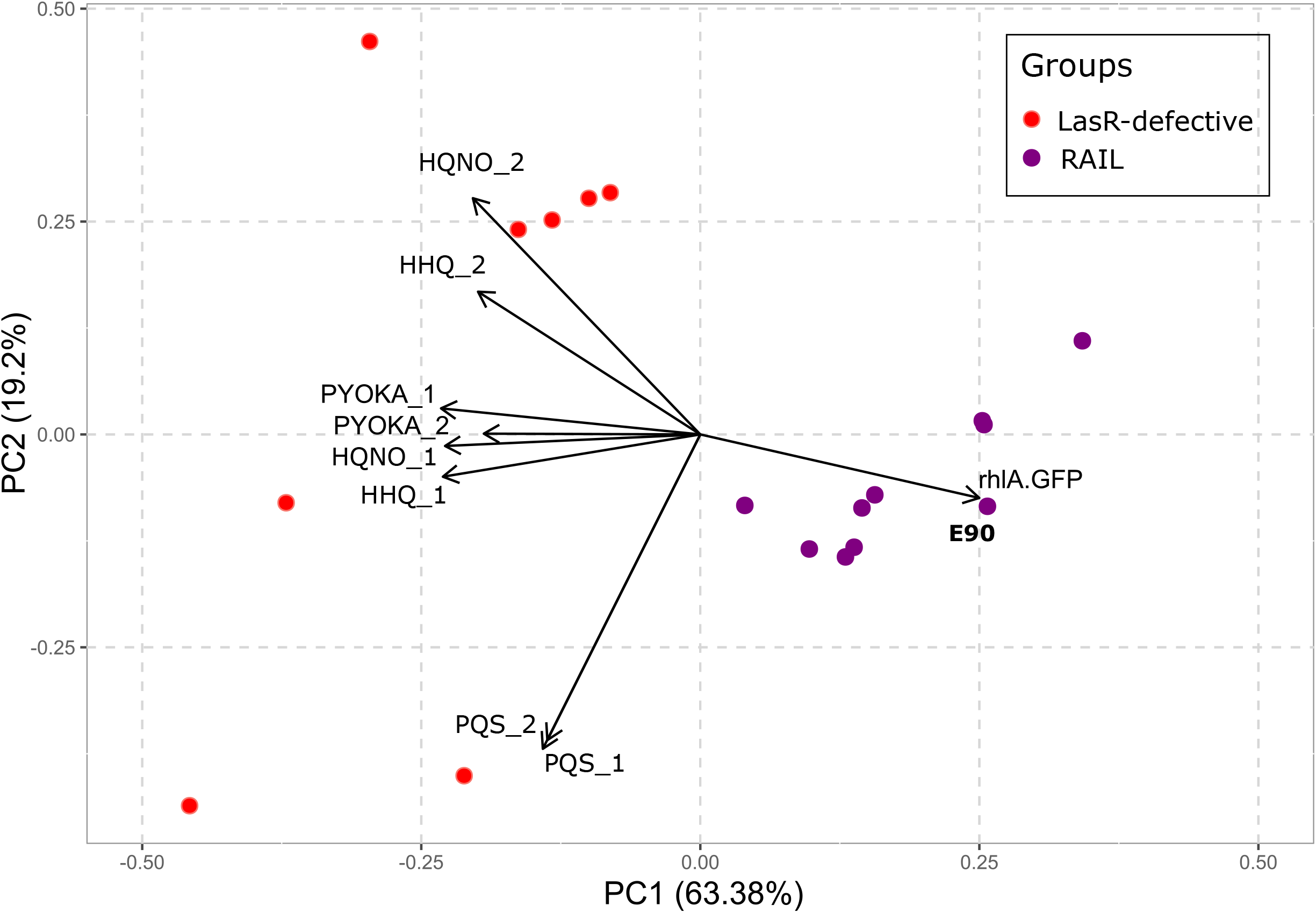
Principal component analyses elaborated with *Pseudomonas aeruginosa* extracellular metabolites analyzed from cultures of LasR-defective isolates. Concentrations of HHQ_1 and 2, HQNO_1 and 2, PQS_1 and 2 sampled at 6 and 24h, respectively. Pyocyanin (PYOKA_1 and PYOKA_2) was sampled at 6 and 24h, respectively. Fluorescence (RFU/OD_600_) from a *rhlA-gfp* reporter was measured at 30h. Reference RhlR Active Independently of LasR (RAIL) strain E90 was included as a control. Groups represent clusters based on a dendrogram elaborated with the same variables (data not shown).

## DISCUSSION

Our knowledge on the prevalence of LasR-defective *P. aeruginosa* isolates is essentially based on clinical samples, mostly from CF or other chronic disease patients (23, 25, 26, 28, 51). It was suggested that adaptation to the CF lungs selects for mutations disturbing LasR activity, resulting in loss of acute virulence factors during chronic infections (26, 52). Mutants in *lasR* were actually associated with CF lung disease progression (25, 53, 54). Still, one study found that LasR-defective isolates were also quite common in severe corneal ulcers, an acute infection, and they were inferred to have an environmental origin (28). Also, subpopulations of LasR-defective mutants have been selected *in vitro* during controlled evolution experiments under conditions that require QS, such as swarming motility, growth as a biofilm and in minimal media with casein as the sole carbon source (55–57) or in a murine chronic wound model (58). Thus, the adaptive pressure for loss of LasR function remains to be clearly defined.

In the present study, we used phenotypic profiling based on the quantification of QS-dependant extracellular molecules to evaluate the prevalence of LasR-defective *P. aeruginosa* strains in environments other than clinical. Isolates were recovered from various environments in Canada and Ivory Coast such as oil-contaminated soils, plumbing systems from hospitals as well as food products from public markets. A LasR-defective strain is one that is deficient in its ability to perform LasR-dependant regulatory activity. Since a LasR-defective profile is not necessarily the consequence of a mutation in the *lasR* gene itself but could also be caused by variations in regulatory elements, we hypothesized that this method would allow for a more accurate and less biased estimation. Indeed, although we did find a good correlation between our classification scheme and the presence of SNPs in the *lasR* gene (**Table 1**), a SNP was found in the sequence of the *lasR* gene of isolate 34JS while this isolate nevertheless has a functional LasR. LasR variations were found in isolates with residual LasR activity in a previous study (23). Our method based on quantifiable phenotypic traits allowed for reliable classification of LasR-defective isolates regardless of the presence of significant modifications in the sequence of the *lasR* gene.

Based on our analyses, a high proportion of environmental isolates are defective in their ability to perform LasR-dependant functions. This is the first time that there is characterisation of such a collection of environmental strains for the prevalence of defective LasR activity. We know only two other studies reporting the presence of a few LasR-defective strains in environmental settings: swimming pools and rivers (27) and dental waterlines units (29). Interestingly, we noted that 23% (9/40) of the strains from the International *Pseudomonas aeruginosa* reference panel are *lasR*- mutants, including three that are not from CF origin (59). Importantly, the percentage of <u>first</u> cultured *P. aeruginosa* isolates from CF patients that are *lasR* mutant is around 20-25% (23, 60, 61). The high prevalence of environmental *lasR-* isolates found here raises a daring question: do these strains really mostly evolve *in vivo* (e.g. CF lungs) as currently thought or could have they been initially acquired by patients directly from the environment?

Very few studies investigated the production profile of HAQs by clinical isolates, but our work shows that it is a relevant indicator of LasR activity. Indeed, LasR is responsible for the activation of the *pqs* operon early during growth but it also controls the transcription of the *pqsH* and *pqsL* genes which encodes for the enzymes responsible for production of PQS and HQNO, respectively. Interestingly, we uncovered the presence of several environmental *P. aeruginosa* isolates defective in the third QS system, mediated by the PQS signal. We did not find variations in the coding sequence of the *pqs* operon or in the *mvfR* (*pqsR*) gene in these strains. Investigations on why these isolates are unable to produce HAQs under our conditions are on-going. Of note, some of these isolates produced significant levels of pyocyanin in King’s A media, which is surprising since a defect in the *pqs* operon should affect the presence of PqsE, a protein essential for full activity of RhlR, the principal activator of pyocyanin production (20). We previously observed a few such strains producing no HAQs but still able to produce pyocyanin, among CF isolates (23). It will be interesting to investigate whether there is an active PqsE in these strains.

RhlR activity is required by *P. aeruginosa* to upregulate functions important for survival in various environments. Our identification of several RAIL strains among our environmental collection highlights the relevance of recent studies where such strains with a rewired QS circuitry have been investigated (22, 23, 50). The expression of RhlR independent of prior LasR activation is forcing us to reconsider the archetypal hierarchy of the *P. aeruginosa* QS cascade. Since full regulation of RhlR-dependant QS depends on PqsE in reference strains such as PA14 (20), it will be interesting to study this interaction in RAIL and HAQ-neg isolates from various environments. The presence of these RhlR-active isolates might explain why some phenotypes such as protease production might not be consistent between LasR-defective strains. For example, many isolates produced no AHL but were still able to grow on casein agar (**Table S2 and S3**).

The prevalence of LasR-deficient isolates was high. While we cannot exclude the possibility that a few isolates coming from the same environment are clonal, which would lead to some over-evaluation, this does not alter our conclusion that LasR-defective strains are neither atypical nor restricted to clinical isolates. Studies on the actual prevalence of LasR-deficient strains depending on the type of environments will be interesting, to uncover the selective pressure for the loss of this regulatory function. Clearly, our understanding *P. aeruginosa*, after all a saprophytic species, is biased by an emphasis on studying clinical isolates. For instance, *P. aeruginosa* also forms Small Colony Variants (SCVs) under specific conditions using phase variation as an adaptation mechanism. This ability was long associated with clinical isolates, but it was found to be conserved among environmental strains as well (62).

Loss of LasR function and HAQ production as well as independent upregulation of RhlR activity appear to be features that are beneficial to *P. aeruginosa* as it adapts to many environmental conditions, not just those of chronic infections. This challenges our traditional view of the QS circuitry of this versatile bacterium, and should be kept in mind when devising approaches to control the virulence and response of this opportunistic pathogen.

## AKNOWLEDGMENTS

The authors would like to thank Sara Habouria and Donatien Comoe Benie for their technical contributions in initial phases of this project. We also thank Marianne Piochon for her expertise with LC/MS analyses. Many thanks to Ajai Dandekar (U. Washington) for providing us with the E90 strain and the *rhlA-gfp* reporter as well as for critical reading of the manuscript and helpful comments.

---

**Figure S1. Clustering of LasR-defective isolates.** A cluster dendrogram was performed on a subset of isolates using the following variables: HHQ_1 and 2, HQNO_1 and 2, PQS_1 and 2 and pyocyanin (PYOKA_1). Concentrations of HHQ_1 and 2, HQNO_1 and 2, PQS_1 and 2 sampled at 6 and 24h. Pyocyanin (PYOKA_2) was sampled at 24h. C4-HSL_1 and 2 and 3-oxo-C12-HSL_1 and 2 were sampled at 3 and 6 h. Values are listed in Table S3. Robustness: **** >90 %, *** 80-89%, ** 70-79%.

**Figure S2. Clustering of LasR-defective isolates.** A cluster dendrogram was performed on 176 isolates using chosen variables HHQ_1 and 2, HQNO_1 and 2, PQS_1 and 2 and pyocyanin (PYOKA_2). Concentrations of HHQ_1 and 2, HQNO_1 and 2, PQS_1 and 2 sampled at 6 and 24h. Pyocyanin (PYOKA_2) was sampled at 24h. Robustness: **** >90 %, *** 80-89%, ** 70-79%.

**Table S1: List of isolates used in this study and their origins.**

**Table S2: Phenotypical characteristics of isolates and reference strains**

**Table S3: AHLs, HHQ, HQNO and PQS and pyocyanin quantifications. All data is concentration in mg/L normalized by total protein concentration (μg/mL) in sample. Expression of *rhlA-gfp* reporter (RFU/OD_600_).**

**Table S4. Whole genome sequence analyses**

## REFERENCES

1. Grosso-Becerra MV, Santos-Medellin C, Gonzalez-Valdez A, Mendez JL, Delgado G, Morales-Espinosa R, et al. *Pseudomonas aeruginosa* clinical and environmental isolates constitute a single population with high phenotypic diversity. BMC Genomics. 2014;15:318.

2. Crone S, Vives-Flórez M, Kvich L, Saunders AM, Malone M, Nicolaisen MH, et al. The environmental occurrence of *Pseudomonas aeruginosa*. APMIS. 2020;128(3):220–31.

3. Abisado RG, Benomar S, Klaus JR, Dandekar AA, Chandler JR. Bacterial Quorum Sensing and Microbial Community Interactions. mBio. 2018;9(3):e02331–17.

4. Gilbert KB, Kim TH, Gupta R, Greenberg EP, Schuster M. Global position analysis of the *Pseudomonas aeruginosa* quorum-sensing transcription factor LasR. Molecular microbiology. 2009;73(6):1072–85.

5. Toder DS, Gambello MJ, Iglewski BH. *Pseudomonas aeruginosa* LasA: a second elastase under the transcriptional control of *lasR*. Molecular microbiology. 1991;5(8):2003–10.

6. Gambello MJ, Iglewski BH. Cloning and characterization of the *Pseudomonas aeruginosa lasR* gene, a transcriptional activator of elastase expression. Journal of bacteriology. 1991;173(9):3000–9.

7. Whiteley M, Lee KM, Greenberg EP. Identification of genes controlled by quorum sensing in *Pseudomonas aeruginosa*. Proc Natl Acad Sci U S A. 1999;96(24):13904–9.

8. Pesci EC, Pearson JP, Seed PC, Iglewski BH. Regulation of *las* and *rhl* quorum sensing in *Pseudomonas aeruginosa*. Journal of bacteriology. 1997;179(10):3127–32.

9. Wagner VE, Bushnell D, Passador L, Brooks AI, Iglewski BH. Microarray analysis of Pseudomonas aeruginosa quorum-sensing regulons: effects of growth phase and environment. Journal of bacteriology. 2003;185(7):2080–95.

10. Latifi A, Foglino M, Tanaka K, Williams P, Lazdunski A. A hierarchical quorum-sensing cascade in *Pseudomonas aeruginosa* links the transcriptional activators LasR and RhIR (VsmR) to expression of the stationary-phase sigma factor RpoS. Molecular Microbiology. 1996;21(6):1137–46.

11. Ochsner UA, Fiechter A, Reiser J. Isolation, characterization, and expression in *Escherichia coli* of the *Pseudomonas aeruginosa rhlAB* genes encoding a rhamnosyltransferase involved in rhamnolipid biosurfactant synthesis. The Journal of biological chemistry. 1994;269(31):19787–95.

12. Brint JM, Ohman DE. Synthesis of multiple exoproducts in *Pseudomonas aeruginosa* is under the control of RhlR-RhlI, another set of regulators in strain PAO1 with homology to the autoinducer-responsive LuxR-LuxI family. Journal of bacteriology. 1995;177(24):7155–63.

13. Albus AM, Pesci EC, Runyen-Janecky LJ, West SE, Iglewski BH. Vfr controls quorum sensing in Pseudomonas aeruginosa. J Bacteriol. 1997;179(12):3928–35.

14. Pessi G, Haas D. Transcriptional control of the hydrogen cyanide biosynthetic genes *hcnABC* by the anaerobic regulator ANR and the quorum-sensing regulators LasR and RhlR in *Pseudomonas aeruginosa*. Journal of bacteriology. 2000;182(24):6940–9.

15. Déziel E, Lépine F, Milot S, He J, Mindrinos MN, Tompkins RG, et al. Analysis of Pseudomonas aeruginosa 4-hydroxy-2-alkylquinolines (HAQs) reveals a role for 4-hydroxy-2-heptylquinoline in cell-to-cell communication. Proceedings of the National Academy of Sciences of the United States of America. 2004;101(5):1339–44.

16. Lightbown JW, Jackson FL. Inhibition of cytochrome systems of heart muscle and certain bacteria by the antagonists of dihydrostreptomycin: 2-alkyl-4-hydroxyquinoline N-oxides. Biochemical Journal. 1956;63(1):130–7.

17. Machan ZA, Taylor GW, Pitt TL, Cole PJ, Wilson R. 2-Heptyl-4-hydroxyquinoline N-oxide, an antistaphylococcal agent produced by *Pseudomonas aeruginosa*. The Journal of antimicrobial chemotherapy. 1992;30(5):615–23.

18. Hazan R, Que YA, Maura D, Strobel B, Majcherczyk PA, Hopper LR, et al. Auto Poisoning of the Respiratory Chain by a Quorum-Sensing-Regulated Molecule Favors Biofilm Formation and Antibiotic Tolerance. Current biology : CB. 2016;26(2):195–206.

19. Mukherjee S, Moustafa DA, Stergioula V, Smith CD, Goldberg JB, Bassler BL. The PqsE and RhlR proteins are an autoinducer synthase-receptor pair that control virulence and biofilm development in *Pseudomonas aeruginosa*. Proceedings of the National Academy of Sciences of the United States of America. 2018;115(40):E9411–E8.

20. Groleau MC, de Oliveira Pereira T, Dekimpe V, Déziel E. PqsE Is Essential for RhlR-Dependent Quorum Sensing Regulation in *Pseudomonas aeruginosa*. mSystems. 2020;5(3).

21. Dekimpe V, Déziel E. Revisiting the quorum-sensing hierarchy in *Pseudomonas aeruginosa*: the transcriptional regulator RhlR regulates LasR-specific factors. Microbiology. 2009;155(Pt 3):712–23.

22. Chen R, Deziel E, Groleau MC, Schaefer AL, Greenberg EP. Social cheating in a *Pseudomonas aeruginosa* quorum-sensing variant. Proc Natl Acad Sci U S A. 2019;116(14):7021–6.

23. Feltner JB, Wolter DJ, Pope CE, Groleau MC, Smalley NE, Greenberg EP, et al. LasR Variant Cystic Fibrosis Isolates Reveal an Adaptable Quorum-Sensing Hierarchy in *Pseudomonas aeruginosa*. mBio. 2016;7(5).

24. Bjarnsholt T, Jensen PO, Jakobsen TH, Phipps R, Nielsen AK, Rybtke MT, et al. Quorum sensing and virulence of *Pseudomonas aeruginosa* during lung infection of cystic fibrosis patients. PLoS One. 2010;5(4):e10115.

25. Hoffman LR, Kulasekara HD, Emerson J, Houston LS, Burns JL, Ramsey BW, et al. *Pseudomonas aeruginosa lasR* mutants are associated with cystic fibrosis lung disease progression. J Cyst Fibros. 2009;8(1):66–70.

26. D’Argenio DA, Wu M, Hoffman LR, Kulasekara HD, Déziel E, Smith EE, et al. Growth phenotypes of *Pseudomonas aeruginosa lasR* mutants adapted to the airways of cystic fibrosis patients. Molecular microbiology. 2007;64(2):512–33.

27. Cabrol S, Olliver A, Pier GB, Andremont A, Ruimy R. Transcription of quorum-sensing system genes in clinical and environmental isolates of *Pseudomonas aeruginosa*. Journal of bacteriology. 2003;185(24):7222–30.

28. Hammond JH, Hebert WP, Naimie A, Ray K, Van Gelder RD, DiGiandomenico A, et al. Environmentally Endemic *Pseudomonas aeruginosa* Strains with Mutations in *lasR* Are Associated with Increased Disease Severity in Corneal Ulcers. mSphere. 2016;1(5).

29. Vincent AT, Freschi L, Jeukens J, Kukavica-Ibrulj I, Emond-Rheault JG, Leduc A, et al. Genomic characterisation of environmental *Pseudomonas aeruginosa* isolated from dental unit waterlines revealed the insertion sequence ISPa11 as a chaotropic element. FEMS microbiology ecology. 2017;93(9).

30. Asfahl KL, Schuster M. Additive Effects of Quorum Sensing Anti-Activators on *Pseudomonas aeruginosa* Virulence Traits and Transcriptome. Frontiers in Microbiology. 2018;8(2654).

31. Lintz MJ, Oinuma K-I, Wysoczynski CL, Greenberg EP, Churchill MEA. Crystal structure of QscR, a *Pseudomonas aeruginosa* quorum sensing signal receptor. Proceedings of the National Academy of Sciences. 2011;108(38):15763–8.

32. Siehnel R, Traxler B, An DD, Parsek MR, Schaefer AL, Singh PK. A unique regulator controls the activation threshold of quorum-regulated genes in *Pseudomonas aeruginosa*. Proceedings of the National Academy of Sciences. 2010;107(17):7916–21.

33. Fan H, Dong Y, Wu D, Bowler MW, Zhang L, Song H. QsIA disrupts LasR dimerization in antiactivation of bacterial quorum sensing. Proceedings of the National Academy of Sciences. 2013;110(51):20765–70.

34. Shah M, Taylor VL, Bona D, Tsao Y, Stanley SY, Pimentel-Elardo SM, et al. A phage-encoded anti-activator inhibits quorum sensing in *Pseudomonas aeruginosa*. Mol Cell. 2021;81(3):571–83 e6.

35. Cabeen MT. Stationary phase-specific virulence factor overproduction by a *lasR* mutant of *Pseudomonas aeruginosa*. PLoS One. 2014;9(2):e88743.

36. Wang Y, Gao L, Rao X, Wang J, Yu H, Jiang J, et al. Characterization of *lasR*-deficient clinical isolates of *Pseudomonas aeruginosa*. Sci Rep. 2018;8(1):13344.

37. Déziel E, Paquette G, Villemur R, Lepine F, Bisaillon J. Biosurfactant production by a soil *pseudomonas* strain growing on polycyclic aromatic hydrocarbons. Applied and environmental microbiology. 1996;62(6):1908–12.

38. Lalancette C, Charron D, Laferriere C, Dolce P, Déziel E, Prevost M, et al. Hospital Drains as Reservoirs of *Pseudomonas aeruginosa*: Multiple-Locus Variable-Number of Tandem Repeats Analysis Genotypes Recovered from Faucets, Sink Surfaces and Patients. Pathogens. 2017;6(3).

39. Benie CK, Dadie A, Guessennd N, N’Gbesso-Kouadio NA, Kouame ND, N’Golo D C, et al. Characterization of Virulence Potential of *Pseudomonas aeruginosa* Isolated from Bovine Meat, Fresh Fish, and Smoked Fish. European journal of microbiology & immunology. 2017;7(1):55–64.

40. Kumar RS, Ayyadurai N, Pandiaraja P, Reddy AV, Venkateswarlu Y, Prakash O, et al. Characterization of antifungal metabolite produced by a new strain *Pseudomonas aeruginosa* PUPa3 that exhibits broad-spectrum antifungal activity and biofertilizing traits. Journal of applied microbiology. 2005;98(1):145–54.

41. Guerra-Santos L, Kappeli O, Fiechter A. *Pseudomonas aeruginosa* biosurfactant production in continuous culture with glucose as carbon source. Applied and environmental microbiology. 1984;48(2):301–5.

42. Rahme LG, Stevens EJ, Wolfort SF, Shao J, Tompkins RG, Ausubel FM. Common virulence factors for bacterial pathogenicity in plants and animals. Science. 1995;268(5219):1899–902.

43. Déziel E, Lépine F, Milot S, He J, Mindrinos MN, Tompkins RG, et al. Analysis of *Pseudomonas aeruginosa* 4-hydroxy-2-alkylquinolines (HAQs) reveals a role for 4-hydroxy-2-heptylquinoline in cell-to-cell communication. Proceedings of the National Academy of Sciences of the United States of America. 2004;101(5):1339–44.

44. King EO, Ward MK, Raney DE. Two simple media for the demonstration of pyocyanin and fluorescin. The Journal of laboratory and clinical medicine. 1954;44(2):301–7.

45. Lépine F, Milot S, Groleau MC, Déziel E. Liquid Chromatography/Mass Spectrometry (LC/MS) for the Detection and Quantification of N-Acyl-L-Homoserine Lactones (AHLs) and 4-Hydroxy-2-Alkylquinolines (HAQs). Methods Mol Biol. 2018;1673:49–59.

46. Team RCD. R: A language and environment for statistical computing. Vienna, Austria2018.

47. Hennig C. Cluster-wise assessment of cluster stability. Computational Statistics & Data Analysis. 2007;52(1):258–71.

48. Wickham H. ggplot2: Elegant Graphics for Data Analysis. New York, NY, USA2009.

49. Freschi L, Jeukens J, Kukavica-Ibrulj I, Boyle B, Dupont MJ, Laroche J, et al. Clinical utilization of genomics data produced by the international *Pseudomonas aeruginosa* consortium. Frontiers in microbiology. 2015;6:1036.

50. Cruz RL, Asfahl KL, Van den Bossche S, Coenye T, Crabbe A, Dandekar AA. RhlR-Regulated Acyl-Homoserine Lactone Quorum Sensing in a Cystic Fibrosis Isolate of *Pseudomonas aeruginosa*. mBio. 2020;11(2).

51. Ciofu O, Mandsberg LF, Bjarnsholt T, Wassermann T, Hoiby N. Genetic adaptation of *Pseudomonas aeruginosa* during chronic lung infection of patients with cystic fibrosis: strong and weak mutators with heterogeneous genetic backgrounds emerge in *mucA* and/or *lasR* mutants. Microbiology (Reading). 2010;156(Pt 4):1108–19.

52. Hoffman LR, Richardson AR, Houston LS, Kulasekara HD, Martens-Habbena W, Klausen M, et al. Nutrient availability as a mechanism for selection of antibiotic tolerant *Pseudomonas aeruginosa* within the CF airway. PLoS pathogens. 2010;6(1):e1000712.

53. Winstanley C, O’Brien S, Brockhurst MA. *Pseudomonas aeruginosa* Evolutionary Adaptation and Diversification in Cystic Fibrosis Chronic Lung Infections. Trends in microbiology. 2016;24(5):327–37.

54. LaFayette SL, Houle D, Beaudoin T, Wojewodka G, Radzioch D, Hoffman LR, et al. Cystic fibrosis-adapted *Pseudomonas aeruginosa* quorum sensing *lasR* mutants cause hyperinflammatory responses. Sci Adv. 2015;1(6).

55. Robitaille S, Groleau MC, Déziel E. Swarming motility growth favours the emergence of a subpopulation of *Pseudomonas aeruginosa* quorum-sensing mutants. Environmental microbiology. 2020;22(7):2892–906.

56. Azimi S, Roberts AEL, Peng S, Weitz JS, McNally A, Brown SP, et al. Allelic polymorphism shapes community function in evolving *Pseudomonas aeruginosa* populations. The ISME journal. 2020;14(8):1929–42.

57. Sandoz KM, Mitzimberg SM, Schuster M. Social cheating in *Pseudomonas aeruginosa* quorum sensing. Proc Natl Acad Sci U S A. 2007;104(40):15876–81.

58. Vanderwoude J, Fleming D, Azimi S, Trivedi U, Rumbaugh KP, Diggle SP. The evolution of virulence in *Pseudomonas aeruginosa* during chronic wound infection. Proceedings Biological sciences. 2020;287(1937):20202272.

59. Freschi L, Bertelli C, Jeukens J, Moore MP, Kukavica-Ibrulj I, Emond-Rheault JG, et al. Genomic characterisation of an international *Pseudomonas aeruginosa* reference panel indicates that the two major groups draw upon distinct mobile gene pools. FEMS microbiology letters. 2018;365(14).

60. Mayer-Hamblett N, Ramsey BW, Kulasekara HD, Wolter DJ, Houston LS, Pope CE, et al. *Pseudomonas aeruginosa* phenotypes associated with eradication failure in children with cystic fibrosis. Clinical infectious diseases : an official publication of the Infectious Diseases Society of America. 2014;59(5):624–31.

61. Mayer-Hamblett N, Rosenfeld M, Gibson RL, Ramsey BW, Kulasekara HD, Retsch-Bogart GZ, et al. *Pseudomonas aeruginosa* in vitro phenotypes distinguish cystic fibrosis infection stages and outcomes. American journal of respiratory and critical care medicine. 2014;190(3):289–97.

62. Besse A, Trottier M, Groleau M-C, Déziel E. The ability of *Pseudomonas aeruginosa* to adopt a Small Colony Variant (SCV) phenotype is conserved, and not restricted to clinical isolates. bioRxiv. 2021:2021.02.05.430018.

