## Supplementary figures and images for "*Pseudomonas aeruginosa* isolates defective in function of the LasR quorum sensing regulator are frequent in diverse environmental niches"

### Supplemental figures

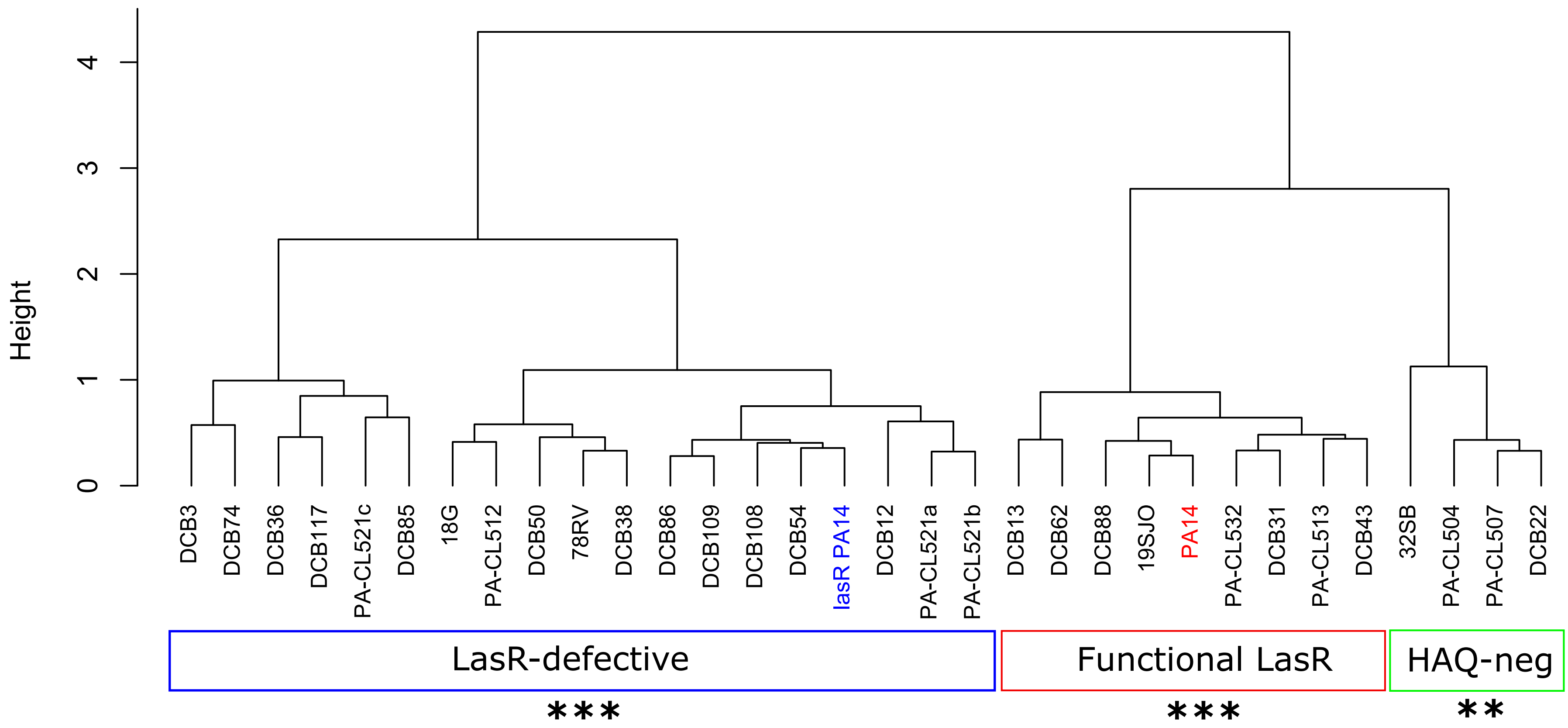

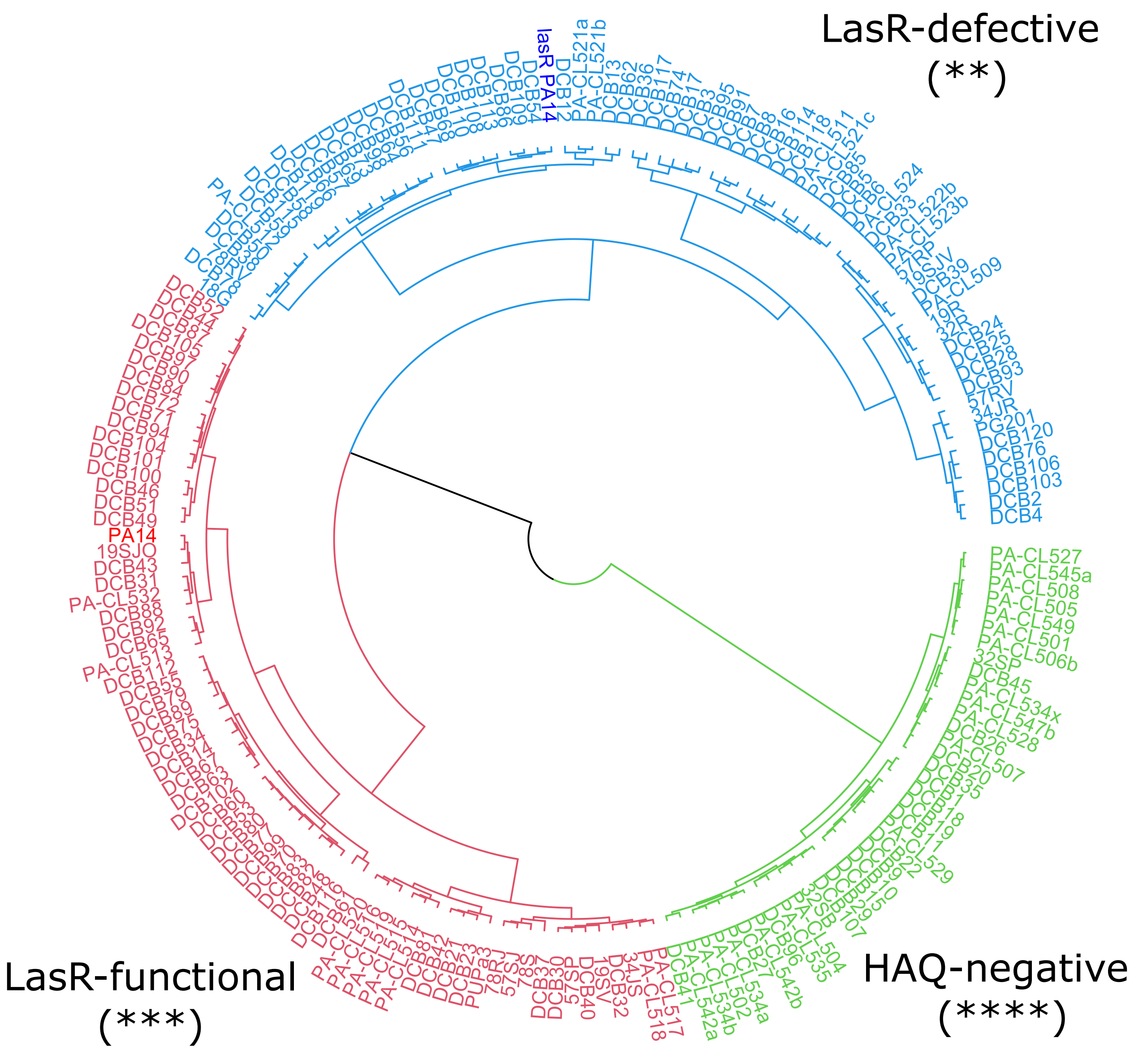
