## Supplemental tables for "*Pseudomonas aeruginosa* isolates defective in function of the LasR quorum sensing regulator are frequent in diverse environmental niches"

**Table S1: List of isolates used in this study and their origin**

| Isolates | Origin | Description | Source |
|---|---|---|---|
| 18G | Thouin's sandpit, Qc, Canada | Contaminated soil (oil residue) | Déziel et al. 1996 |
| 19SJO | Thouin's sandpit, Qc, Canada | Contaminated soil (oil residue) | Déziel et al. 1996 |
| 19SJV | Thouin's sandpit, Qc, Canada | Contaminated soil (oil residue) | Déziel et al. 1996 |
| 19R | Thouin's sandpit, Qc, Canada | Contaminated soil (oil residue) | Déziel et al. 1996 |
| 19SV | Thouin's sandpit, Qc, Canada | Contaminated soil (oil residue) | Déziel et al. 1996 |
| 32SP | Thouin's sandpit, Qc, Canada | Contaminated soil (oil residue) | Déziel et al. 1996 |
| 32SB | Thouin's sandpit, Qc, Canada | Contaminated soil (oil residue) | Déziel et al. 1996 |
| 32R | Thouin's sandpit, Qc, Canada | Contaminated soil (oil residue) | Déziel et al. 1996 |
| 34JS | Thouin's sandpit, Qc, Canada | Contaminated soil (oil residue) | Déziel et al. 1996 |
| 34JR | Thouin's sandpit, Qc, Canada | Contaminated soil (oil residue) | Déziel et al. 1996 |
| 57SJ | Thouin's sandpit, Qc, Canada | Contaminated soil (oil residue) | Déziel et al. 1996 |
| 57SP | Thouin's sandpit, Qc, Canada | Contaminated soil (oil residue) | Déziel et al. 1996 |
| 57RV | Thouin's sandpit, Qc, Canada | Contaminated soil (oil residue) | Déziel et al. 1996 |
| 57RP | Thouin's sandpit, Qc, Canada | Contaminated soil (oil residue) | Déziel et al. 1996 |
| 78RJ | Thouin's sandpit, Qc, Canada | Contaminated soil (oil residue) | Déziel et al. 1996 |
| 78RV | Thouin's sandpit, Qc, Canada | Contaminated soil (oil residue) | Déziel et al. 1996 |
| 78S | Thouin's sandpit, Qc, Canada | Contaminated soil (oil residue) | Déziel et al. 1996 |
| PG201 | Zurich, Switzerland | Soil | Guerra-Santos et al. 1984 |
| PUPa3 | Rice rhizosphere, India | Soil | Kumar et al. 2005 |
| PA-CL501 | Hospital sink, Montreal, Canada | Drain | Lalancette et al. 2017 |
| PA-CL502 | Hospital sink, Montreal, Canada | Drain | Lalancette et al. 2017 |
| PA-CL504 | Hospital sink, Montreal, Canada | Drain | Lalancette et al. 2017 |
| PA-CL505 | Hospital sink, Montreal, Canada | Drain | Lalancette et al. 2017 |
| PA-CL506b | Hospital sink, Montreal, Canada | Drain | Lalancette et al. 2017 |
| PA-CL507 | Hospital sink, Montreal, Canada | Drain | Lalancette et al. 2017 |
| PA-CL508 | Hospital sink, Montreal, Canada | Drain | Lalancette et al. 2017 |
| PA-CL509 | Hospital sink, Montreal, Canada | Drain | Lalancette et al. 2017 |
| PA-CL511 | Hospital sink, Montreal, Canada | Drain | Lalancette et al. 2017 |
| PA-CL512 | Hospital sink, Montreal, Canada | Drain | Lalancette et al. 2017 |
| PA-CL513 | Hospital sink, Montreal, Canada | Drain | Lalancette et al. 2017 |
| PA-CL514 | Hospital sink, Montreal, Canada | Drain | Lalancette et al. 2017 |
| PA-CL515 | Hospital sink, Montreal, Canada | Drain | Lalancette et al. 2017 |
| PA-CL516 | Hospital sink, Montreal, Canada | Drain | Lalancette et al. 2017 |
| PA-CL517 | Hospital sink, Montreal, Canada | Drain | Lalancette et al. 2017 |
| PA-CL518 | Hospital sink, Montreal, Canada | Drain | Lalancette et al. 2017 |
| PA-CL519 | Hospital sink, Montreal, Canada | Drain | Lalancette et al. 2017 |
| PA-CL520 | Hospital sink, Montreal, Canada | Drain | Lalancette et al. 2017 |
| PA-CL521a | Hospital sink, Montreal, Canada | Drain | Lalancette et al. 2017 |
| PA-CL521b | Hospital sink, Montreal, Canada | Drain | Lalancette et al. 2017 |
| PA-CL521c | Hospital sink, Montreal, Canada | Drain | Lalancette et al. 2017 |
| PA-CL522b | Hospital sink, Montreal, Canada | Drain | Lalancette et al. 2017 |
| PA-CL523b | Hospital sink, Montreal, Canada | Splash | Lalancette et al. 2017 |
| PA-CL524 | Hospital sink, Montreal, Canada | Splash | Lalancette et al. 2017 |
| PA-CL527 | Hospital sink, Montreal, Canada | Drain | Lalancette et al. 2017 |

| | | | |
|---|---|---|---|
| PA-CL528 | Hospital sink, Montreal, Canada | Drain | Lalancette et al. 2017 |
| PA-CL529 | Hospital sink, Montreal, Canada | Drain | Lalancette et al. 2017 |
| PA-CL532 | Hospital sink, Montreal, Canada | Splash | Lalancette et al. 2017 |
| PA-CL534a | Hospital sink, Montreal, Canada | Faucet | Lalancette et al. 2017 |
| PA-CL534b | Hospital sink, Montreal, Canada | Faucet | Lalancette et al. 2017 |
| PA-CL534x | Hospital sink, Montreal, Canada | Faucet | Lalancette et al. 2017 |
| PA-CL535 | Hospital sink, Montreal, Canada | Splash | Lalancette et al. 2017 |
| PA-CL542a | Hospital sink, Montreal, Canada | Splash | Lalancette et al. 2017 |
| PA-CL542b | Hospital sink, Montreal, Canada | Splash | Lalancette et al. 2017 |
| PA-CL545a | Hospital sink, Montreal, Canada | Faucet | Lalancette et al. 2017 |
| PA-CL547b | Hospital sink, Montreal, Canada | Faucet | Lalancette et al. 2017 |
| PA-CL549 | Hospital sink, Montreal, Canada | Splash | Lalancette et al. 2017 |
| DCB1; YOP VB5 C1 | Abidjan, Ivory Coast | Bovine meat | Benie et al., 2017 |
| DCB2; ABO VB 29 C1 | Abidjan, Ivory Coast | Bovine meat | Benie et al., 2017 |
| DCB3; ABO PFR11 C1 | Abidjan, Ivory Coast | Fresh fish | Benie et al., 2017 |
| DCB4; YOP VB24 C1 | Abidjan, Ivory Coast | Bovine meat | Benie et al., 2017 |
| DCB5; ABO VB 48 C1 | Abidjan, Ivory Coast | Bovine meat | Benie et al., 2017 |
| DCB6; BG VB5 C2 | Abidjan, Ivory Coast | Bovine meat | Benie et al., 2017 |
| DCB7; ABO VB50 C1 | Abidjan, Ivory Coast | Bovine meat | Benie et al., 2017 |
| DCB8; PB VB19 C1 | Abidjan, Ivory Coast | Bovine meat | Benie et al., 2017 |
| DCB9; PB VB6 C1 | Abidjan, Ivory Coast | Bovine meat | Benie et al., 2017 |
| DCB10; BG VB5 C | Abidjan, Ivory Coast | Bovine meat | Benie et al., 2017 |
| DCB11; PB VB13 C1 | Abidjan, Ivory Coast | Bovine meat | Benie et al., 2017 |
| DCB12; ABO VB30 C1 | Abidjan, Ivory Coast | Bovine meat | Benie et al., 2017 |
| DCB13; YOP VB2 C3 | Abidjan, Ivory Coast | Bovine meat | Benie et al., 2017 |
| DCB14; PB VB16 C3 | Abidjan, Ivory Coast | Bovine meat | Benie et al., 2017 |
| DCB15; ADJ VB5 C1 | Abidjan, Ivory Coast | Bovine meat | Benie et al., 2017 |
| DCB16; ADJ VB2 C1 | Abidjan, Ivory Coast | Bovine meat | Benie et al., 2017 |
| DCB17; PB VB15 C1 | Abidjan, Ivory Coast | Bovine meat | Benie et al., 2017 |
| DCB18; YOP PFR12 C3 | Abidjan, Ivory Coast | Fresh fish | Benie et al., 2017 |
| DCB19; BG VB11 C1 | Abidjan, Ivory Coast | Bovine meat | Benie et al., 2017 |
| DCB20; ADJ PF1 C1 | Abidjan, Ivory Coast | Smoked fish | Benie et al., 2017 |
| DCB21; BG VB1 C2 | Abidjan, Ivory Coast | Bovine meat | Benie et al., 2017 |
| DCB22; ABO PFR5 C2 | Abidjan, Ivory Coast | Fresh fish | Benie et al., 2017 |
| DCB23; ABO VB7 C1 | Abidjan, Ivory Coast | Bovine meat | Benie et al., 2017 |
| DCB24; BG VB1 C3 | Abidjan, Ivory Coast | Bovine meat | Benie et al., 2017 |
| DCB25; ABO VB52 C1 | Abidjan, Ivory Coast | Bovine meat | Benie et al., 2017 |
| DCB26; ADJ VB7 C1 | Abidjan, Ivory Coast | Bovine meat | Benie et al., 2017 |
| DCB27; ABO VB3 C1 | Abidjan, Ivory Coast | Bovine meat | Benie et al., 2017 |
| DCB28; YOP VB2 C2 | Abidjan, Ivory Coast | Bovine meat | Benie et al., 2017 |
| DCB29; PB PFR5 C1 | Abidjan, Ivory Coast | Fresh fish | Benie et al., 2017 |
| DCB30; ABO PF3 C3 | Abidjan, Ivory Coast | Smoked fish | Benie et al., 2017 |
| DCB31; ABO PF5 C1 | Abidjan, Ivory Coast | Smoked fish | Benie et al., 2017 |
| DCB32; BG PFR4 | Abidjan, Ivory Coast | Smoked fish | Benie et al., 2017 |
| DCB33; ABO PFR9 | Abidjan, Ivory Coast | Fresh fish | Benie et al., 2017 |
| DCB34; PB PFR11 C2 | Abidjan, Ivory Coast | Fresh fish | Benie et al., 2017 |
| DCB35; PB PFR1 C1 | Abidjan, Ivory Coast | Smoked fish | Benie et al., 2017 |

| | | | |
|---|---|---|---|
| DCB36; ABO VB5 C2 | Abidjan, Ivory Coast | Bovine meat | Benie et al., 2017 |
| DCB37; ABO VB3 C2 | Abidjan, Ivory Coast | Bovine meat | Benie et al., 2017 |
| DCB38; ABO VB5 C3 | Abidjan, Ivory Coast | Bovine meat | Benie et al., 2017 |
| DCB39; PB PFR12 C1 | Abidjan, Ivory Coast | Fresh fish | Benie et al., 2017 |
| DCB40; ABO PF5 C3 | Abidjan, Ivory Coast | Smoked fish | Benie et al., 2017 |
| DCB41; ABO PFR1 C2 | Abidjan, Ivory Coast | Fresh fish | Benie et al., 2017 |
| DCB42; ABO VB44 C1 | Abidjan, Ivory Coast | Bovine meat | Benie et al., 2017 |
| DCB43; ADJ PF14 C2 | Abidjan, Ivory Coast | Smoked fish | Benie et al., 2017 |
| DCB44; PB PFR5 C2 | Abidjan, Ivory Coast | Fresh fish | Benie et al., 2017 |
| DCB45; ABO PF10 C1 | Abidjan, Ivory Coast | Smoked fish | Benie et al., 2017 |
| DCB46; ABO PFR1 C1 | Abidjan, Ivory Coast | Fresh fish | Benie et al., 2017 |
| DCB47; YOP PFR15 C2 | Abidjan, Ivory Coast | Fresh fish | Benie et al., 2017 |
| DCB48; ABO PF12 | Abidjan, Ivory Coast | Smoked fish | Benie et al., 2017 |
| DCB49; PB VB7 C3 | Abidjan, Ivory Coast | Bovine meat | Benie et al., 2017 |
| DCB50; ABO PF26 | Abidjan, Ivory Coast | Smoked fish | Benie et al., 2017 |
| DCB51; BG PFR9 | Abidjan, Ivory Coast | Fresh fish | Benie et al., 2017 |
| DCB52; ADJ PFR19 | Abidjan, Ivory Coast | Fresh fish | Benie et al., 2017 |
| DCB53; ADJ VB5 C3 | Abidjan, Ivory Coast | Bovine meat | Benie et al., 2017 |
| DCB54; ABO PF3 C2 | Abidjan, Ivory Coast | Smoked fish | Benie et al., 2017 |
| DCB55; BG VB2 C2 | Abidjan, Ivory Coast | Bovine meat | Benie et al., 2017 |
| DCB56; ABO VB7 C2 | Abidjan, Ivory Coast | Bovine meat | Benie et al., 2017 |
| DCB57; YOP VB15 C1 | Abidjan, Ivory Coast | Bovine meat | Benie et al., 2017 |
| DCB58; YOP VB5 C3 | Abidjan, Ivory Coast | Bovine meat | Benie et al., 2017 |
| DCB59; PB VB22 C1 | Abidjan, Ivory Coast | Bovine meat | Benie et al., 2017 |
| DCB60; PB VB19 C2 | Abidjan, Ivory Coast | Bovine meat | Benie et al., 2017 |
| DCB61; PB VB6 C2 | Abidjan, Ivory Coast | Bovine meat | Benie et al., 2017 |
| DCB62; PB VB17 C1 | Abidjan, Ivory Coast | Bovine meat | Benie et al., 2017 |
| DCB63; ADJ PFR17 | Abidjan, Ivory Coast | Fresh fish | Benie et al., 2017 |
| DCB64; YOP VB13 C1 | Abidjan, Ivory Coast | Bovine meat | Benie et al., 2017 |
| DCB65; ADJ PFR21 | Abidjan, Ivory Coast | Fresh fish | Benie et al., 2017 |
| DCB66; ABO VB6 C3 | Abidjan, Ivory Coast | Bovine meat | Benie et al., 2017 |
| DCB67; PB VB8 C1 | Abidjan, Ivory Coast | Bovine meat | Benie et al., 2017 |
| DCB68; PB VB22 C1 | Abidjan, Ivory Coast | Bovine meat | Benie et al., 2017 |
| DCB69; ADJ PF23 | Abidjan, Ivory Coast | Smoked fish | Benie et al., 2017 |
| DCB70; YOP PFR12 C3 | Abidjan, Ivory Coast | Fresh fish | Benie et al., 2017 |
| DCB71; ADJ PFR12 | Abidjan, Ivory Coast | Fresh fish | Benie et al., 2017 |
| DCB72; ADJ PFR16 | Abidjan, Ivory Coast | Fresh fish | Benie et al., 2017 |
| DCB73; ADJ PFR15 | Abidjan, Ivory Coast | Fresh fish | Benie et al., 2017 |
| DCB74; PB VB14 C1 | Abidjan, Ivory Coast | Bovine meat | Benie et al., 2017 |
| DCB75; ADJ PF20 C1 | Abidjan, Ivory Coast | Smoked fish | Benie et al., 2017 |
| DCB76; ABO VB6 C1 | Abidjan, Ivory Coast | Bovine meat | Benie et al., 2017 |
| DCB77; PB VB9 C3 | Abidjan, Ivory Coast | Bovine meat | Benie et al., 2017 |
| DCB78; PB VB8 C2 | Abidjan, Ivory Coast | Bovine meat | Benie et al., 2017 |
| DCB79; PB PFR7 C1 | Abidjan, Ivory Coast | Fresh fish | Benie et al., 2017 |
| DCB80; PBVB2 C2 | Abidjan, Ivory Coast | Bovine meat | Benie et al., 2017 |
| DCB81; PB VB7 C1 | Abidjan, Ivory Coast | Bovine meat | Benie et al., 2017 |
| DCB82; ABO VB4 C2 | Abidjan, Ivory Coast | Bovine meat | Benie et al., 2017 |

| | | | |
|---|---|---|---|
| DCB83; ADJ PFR10 C1 | Abidjan, Ivory Coast | Fresh fish | Benie et al., 2017 |
| DCB84; PB VB13 C2 | Abidjan, Ivory Coast | Bovine meat | Benie et al., 2017 |
| DCB85; ADJ VB12 C1 | Abidjan, Ivory Coast | Bovine meat | Benie et al., 2017 |
| DCB86; ABO VB4C1 | Abidjan, Ivory Coast | Bovine meat | Benie et al., 2017 |
| DCB87; ABO PF11 C1 | Abidjan, Ivory Coast | Smoked fish | Benie et al., 2017 |
| DCB88; ADJ PFR26 C1 | Abidjan, Ivory Coast | Fresh fish | Benie et al., 2017 |
| DCB89; ADJ VB3 C2 | Abidjan, Ivory Coast | Bovine meat | Benie et al., 2017 |
| DCB90; ADJ PFR4 C1 | Abidjan, Ivory Coast | Fresh fish | Benie et al., 2017 |
| DCB91; YOP PFR2 C1 | Abidjan, Ivory Coast | Fresh fish | Benie et al., 2017 |
| DCB92; ABO PF15 | Abidjan, Ivory Coast | Smoked fish | Benie et al., 2017 |
| DCB93; ABO VB3 C3 | Abidjan, Ivory Coast | Bovine meat | Benie et al., 2017 |
| DCB94; PB VB9 C2 | Abidjan, Ivory Coast | Bovine meat | Benie et al., 2017 |
| DCB95; ADJ PF14 C1 | Abidjan, Ivory Coast | Smoked fish | Benie et al., 2017 |
| DCB96; ADJ PF9C1 | Abidjan, Ivory Coast | Smoked fish | Benie et al., 2017 |
| DCB97; ADJ PF6C1 | Abidjan, Ivory Coast | Smoked fish | Benie et al., 2017 |
| DCB98; ADJ PF13 | Abidjan, Ivory Coast | Smoked fish | Benie et al., 2017 |
| DCB99; YOP PF29C1 | Abidjan, Ivory Coast | Smoked fish | Benie et al., 2017 |
| DCB100; YOP PFR12 C2 | Abidjan, Ivory Coast | Fresh fish | Benie et al., 2017 |
| DCB101; ENVABO | Abidjan, Ivory Coast | Environment | Benie et al., 2017 |
| DCB102; ENVABO | Abidjan, Ivory Coast | Environment | Benie et al., 2017 |
| DCB103; ENVABO | Abidjan, Ivory Coast | Environment | Benie et al., 2017 |
| DCB104; ENVABO | Abidjan, Ivory Coast | Environment | Benie et al., 2017 |
| DCB105; ENVABO | Abidjan, Ivory Coast | Environment | Benie et al., 2017 |
| DCB106; ENVADJ | Abidjan, Ivory Coast | Environment | Benie et al., 2017 |
| DCB107; ENVADJ | Abidjan, Ivory Coast | Environment | Benie et al., 2017 |
| DCB108; ENVADJ | Abidjan, Ivory Coast | Environment | Benie et al., 2017 |
| DCB109; ENVADJ | Abidjan, Ivory Coast | Environment | Benie et al., 2017 |
| DCB110; ENVADJ | Abidjan, Ivory Coast | Environment | Benie et al., 2017 |
| DCB111; ENVYOP | Abidjan, Ivory Coast | Environment | Benie et al., 2017 |
| DCB112; ENVYOP | Abidjan, Ivory Coast | Environment | Benie et al., 2017 |
| DCB113; ENVYOP | Abidjan, Ivory Coast | Environment | Benie et al., 2017 |
| DCB114; ENVYOP | Abidjan, Ivory Coast | Environment | Benie et al., 2017 |
| DCB115; ENVYOP | Abidjan, Ivory Coast | Environment | Benie et al., 2017 |
| DCB116; ENVPBT | Abidjan, Ivory Coast | Environment | Benie et al., 2017 |
| DCB117; ENVPBT | Abidjan, Ivory Coast | Environment | Benie et al., 2017 |
| DCB118; ENVPBT | Abidjan, Ivory Coast | Environment | Benie et al., 2017 |
| DCB119; ENVBG | Abidjan, Ivory Coast | Environment | Benie et al., 2017 |
| DCB120; ENVBG | Abidjan, Ivory Coast | Environment | Benie et al., 2017 |

**Table S2: Phenotypical caracteristics of isolates and reference strains**

| Isolates | Autolysis | Metallic sheen | PYO TSB | PYO King's A | Protease production[1] |
|---|---|---|---|---|---|
| PA14 | - | - | + | - | + |
| PA14 *lasR* ::Gm | + | + | - | + | - |
| 18G | + | + | - | + | - |
| 19SJO | - | - | - | - | + |
| 19SJV | + | + | - | + | - |
| 19R | + | + | - | + | - |
| 19SV | - | - | - | + | + |
| 32SP | - | - | - | + | - |
| 32SB | - | - | - | + | - |
| 32R | + | + | - | + | - |
| 34JS | + | - | - | + | + |
| 34JR | + | + | - | + | - |
| 57SJ | - | - | - | + | + |
| 57SP | + | +, - | - | + | + |
| 57RV | + | + | - | + | - |
| 57RP | + | + | - | + | - |
| 78RJ | + | + | - | + | + |
| 78RV | - | + | - | + | - |
| 78S | + | - | - | + | + |
| PG201 | - | - | - | + | - |
| PUPa3 | - | - | - | - | - |
| PA-CL501 | - | - | - | + | + |
| PA-CL502 | - | - | - | - | + |
| PA-CL504 | - | - | - | + | + |
| PA-CL505 | - | - | - | + | + |
| PA-CL506b | - | - | - | + | + |
| PA-CL507 | - | + | - | + | + |
| PA-CL508 | - | - | - | + | - |
| PA-CL509 | + | + | - | + | - |
| PA-CL511 | + | + | - | + | - |
| PA-CL512 | + | + | - | + | + |
| PA-CL513 | - | - | + | + | + |
| PA-CL514 | - | - | - | - | + |
| PA-CL515 | - | - | - | - | + |
| PA-CL516 | - | - | - | - | + |
| PA-CL517 | - | - | - | + | + |
| PA-CL518 | - | - | - | - | + |
| PA-CL519 | - | - | - | + | + |
| PA-CL520 | - | - | - | + | + |
| PA-CL521a | + | - | - | + | + |

| | | | | | |
|---|---|---|---|---|---|
| PA-CL521b | + | + | - | + | + |
| PA-CL521c | - | - | - | - | - |
| PA-CL522b | + | - | - | + | - |
| PA-CL523b | - | +, - | - | + | + |
| PA-CL524 | - | - | - | + | - |
| PA-CL527 | - | - | - | - | + |
| PA-CL528 | - | - | - | + | + |
| PA-CL529 | - | - | - | - | + |
| PA-CL532 | - | - | + | + | + |
| PA-CL534a | - | - | - | + | + |
| PA-CL534b | - | - | - | - | + |
| PA-CL534x | - | - | - | - | + |
| PA-CL535 | - | - | - | - | + |
| PA-CL542a | - | - | - | - | + |
| PA-CL542b | - | - | - | - | + |
| PA-CL545a | - | - | - | - | + |
| PA-CL547b | - | - | - | - | + |
| PA-CL549 | - | - | - | - | + |
| DCB1; YOP VB5 C1 | - | - | - | + | + |
| DCB2; ABO VB 29 C1 | + | - | - | + | + |
| DCB3; ABO PFR11 C1 | +, - | - | - | - | + |
| DCB4; YOP VB24 C1 | - | - | - | + | + |
| DCB5; ABO VB 48 C1 | - | - | - | - | + |
| DCB6; BG VB5 C2 | - | - | - | + | + |
| DCB7; ABO VB50 C1 | + | - | - | + | + |
| DCB8; PB VB19 C1 | - | - | - | + | + |
| DCB9; PB VB6 C1 | - | - | - | + | + |
| DCB10; BG VB5 C | - | - | - | + | + |
| DCB11; PB VB13 C1 | - | - | - | - | + |
| DCB12; ABO VB30 C1 | - | - | - | + | + |
| DCB13; YOP VB2 C3 | - | - | + | + | + |
| DCB14; PB VB16 C3 | - | - | - | + | + |
| DCB15; ADJ VB5 C1 | - | - | - | + | + |
| DCB16; ADJ VB2 C1 | - | - | - | + | + |
| DCB17; PB VB15 C1 | +, - | - | - | - | + |
| DCB18; YOP PFR12 C3 | - | - | - | + | + |
| DCB19; BG VB11 C1 | - | - | - | - | + |
| DCB20; ADJ PF1 C1 | - | - | - | + | + |
| DCB21; BG VB1 C2 | - | - | + | - | + |
| DCB22; ABO PFR5 C2 | - | - | - | - | + |
| DCB23; ABO VB7 C1 | - | - | + | + | + |
| DCB24; BG VB1 C3 | - | - | - | + | + |
| DCB25; ABO VB52 C1 | - | - | - | + | + |
| DCB26; ADJ VB7 C1 | - | - | - | - | + |
| DCB27; ABO VB3 C1 | - | - | - | - | + |

| | | | | | |
|---|---|---|---|---|---|
| DCB28; YOP VB2 C2 | + + | + | + | + | + |
| DCB29; PB PFR5 C1 | - | - | - | + | + |
| DCB30; ABO PF3 C3 | - | - | - | - | + |
| DCB31; ABO PF5 C1 | - | - | - | - | + |
| DCB32; BG PFR4 | - | - | - | + | + |
| DCB33; ABO PFR9 | + | + | - | + | + |
| DCB34; PB PFR11 C2 | - | - | + | + | + |
| DCB35; PB PFR1 C1 | - | - | - | - | + |
| DCB36; ABO VB5 C2 | - | - | - | + | + |
| DCB37; ABO VB3 C2 | - | - | - | - | + |
| DCB38; ABO VB5 C3 | + + + | + | - | + | + |
| DCB39; PB PFR12 C1 | + + | + + | - | + | + |
| DCB40; ABO PF5 C3 | - | - | - | - | + |
| DCB41; ABO PFR1 C2 | - | - | - | - | + |
| DCB42; ABO VB44 C1 | - | - | - | - | + |
| DCB43; ADJ PF14 C2 | - | - | - | + | + |
| DCB44; PB PFR5 C2 | - | - | - | + | + |
| DCB45; ABO PF10 C1 | - | - | - | + | + |
| DCB46; ABO PFR1 C1 | - | - | - | + | + |
| DCB47; YOP PFR15 C2 | - | - | - | + | + |
| DCB48; ABO PF12 | - | - | - | + | + |
| DCB49; PB VB7 C3 | - | - | - | + | + |
| DCB50; ABO PF26 | + | + | - | + | + |
| DCB51; BG PFR9 | - | - | - | - | + |
| DCB52; ADJ PFR19 | - | - | - | - | + |
| DCB53; ADJ VB5 C3 | - | - | - | + | + |
| DCB54; ABO PF3 C2 | + + | + + | - | + | + |
| DCB55; BG VB2 C2 | + + | - | - | + | + |
| DCB56; ABO VB7 C2 | + + | - | - | + | + |
| DCB57; YOP VB15 C1 | + + | + + | - | + | + |
| DCB58; YOP VB5 C3 | + + | + + | - | + | + |
| DCB59; PB VB22 C1 | + + | + + | - | + | + |
| DCB60; PB VB19 C2 | - | - | - | + | + |
| DCB61; PB VB6 C2 | - | - | + | + | + |
| DCB62; PB VB17 C1 | + + | + + | + | + | + |
| DCB63; ADJ PFR17 | + + | + + | + | + | + |
| DCB64; YOP VB13 C1 | + + | + | - | + | + |
| DCB65; ADJ PFR21 | - | - | - | + | + |
| DCB66; ABO VB6 C3 | - | - | - | + | + |
| DCB67; PB VB8 C1 | + + + | + + + | - | + | + |
| DCB68; PB VB22 C1 | + + + | + + + | - | + | + |
| DCB69; ADJ PF23 | + + + | + | - | + | + |
| DCB70; YOP PFR12 C3 | + + | +, - | - | + | + |
| DCB71; ADJ PFR12 | - | - | + | + | + |
| DCB72; ADJ PFR16 | - | - | + | + | + |

| | | | | | |
|---|---|---|---|---|---|
| DCB73; ADJ PFR15 | + | - | - | + | + |
| DCB74; PB VB14 C1 | - | - | - | - | + |
| DCB75; ADJ PF20 C1 | + + + | + | + | + | + |
| DCB76; ABO VB6 C1 | - | - | - | + | + |
| DCB77; PB VB9 C3 | - | - | - | + | + |
| DCB78; PB VB8 C2 | +, - | - | - | - | + |
| DCB79; PB PFR7 C1 | - | - | - | + | + |
| DCB80; PBVB2 C2 | - | - | - | + | + |
| DCB81; PB VB7 C1 | - | - | - | - | + |
| DCB82; ABO VB4 C2 | +, - | - | - | + | + |
| DCB83; ADJ PFR10 C1 | - | - | + | + | + |
| DCB84; PB VB13 C2 | - | - | + | + | - |
| DCB85; ADJ VB12 C1 | - | - | - | + | + |
| DCB86; ABO VB4C1 | +, - | - | - | + | + |
| DCB87; ABO PF11 C1 | - | - | - | + | + |
| DCB88; ADJ PFR26 C1 | - | - | - | - | + |
| DCB89; ADJ VB3 C2 | - | - | - | + | + |
| DCB90; ADJ PFR4 C1 | - | - | - | - | - |
| DCB91; YOP PFR2 C1 | +, - | +, - | - | + | + |
| DCB92; ABO PF15 | +, - | - | + | + | + |
| DCB93; ABO VB3 C3 | - | - | - | - | + |
| DCB94; PB VB9 C2 | - | - | - | + | - |
| DCB95; ADJ PF14 C1 | +, - | - | - | - | + |
| DCB96; ADJ PF9C1 | - | - | - | - | + |
| DCB97; ADJ PF6C1 | - | - | + | + | + |
| DCB98; ADJ PF13 | - | - | - | + | + |
| DCB99; YOP PF29C1 | - | - | + | + | + |
| DCB100; YOP PFR12 C[illegible] | - | - | + | + | + |
| DCB101; ENVABO | - | - | + | - | + |
| DCB102; ENVABO | + | - | - | + | +, - |
| DCB103; ENVABO | - | - | - | - | + |
| DCB104; ENVABO | - | - | + | + | + |
| DCB105; ENVABO | - | - | + | + | + |
| DCB106; ENVADJ | - | - | - | + | + |
| DCB107; ENVADJ | - | - | - | + | + |
| DCB108; ENVADJ | + | +, - | - | + | + |
| DCB109; ENVADJ | - | - | - | + | + |
| DCB110; ENVADJ | +, - | - | - | + | + |
| DCB111; ENVYOP | - | - | - | + | + |
| DCB112; ENVYOP | - | - | - | + | + |
| DCB113; ENVYOP | - | - | - | + | + |
| DCB114; ENVYOP | - | - | - | + | + |
| DCB115; ENVYOP | - | - | - | - | - |
| DCB116; ENVPBT | + | + | - | + | +, - |
| DCB117; ENVPBT | - | - | - | + | + |

| | | | | | |
|---|---|---|---|---|---|
| DCB118; ENVPBT | - | - | - | + | + |
| DCB119; ENVBG | - | - | - | + | + |
| DCB120; ENVBG | - | - | - | - | + |
| E90 | + | + | - | + | + |

[1] Growth on casein agar

+ : presence of

- : absence of

**Table S3 Concentrations of AHLs, HHQ, HQNO and PQS and pyocyanin quantification. All data is normalized by total protein concentration in sample. Expression of rhlA-gfp reporter (RFU/OD600).**

| | 3-oxo-C12-HSL_1 | 3-oxo-C12-HSL_2 | C4-HSL_1 | C4-HSL_2 | PYOKA_1 | PYOKA_2 | HHQ_1 | HHQ_2 | HQNO_1 | HQNO_2 | PQS_1 | PQS_1 | rhlA-gfp | Final classification |
|---|---|---|---|---|---|---|---|---|---|---|---|---|---|---|
| 18G | 0.000 | 0.000 | 0.000 | 0.001 | 0.022 | 0.028 | 0.148 | 0.197 | 0.027 | 0.092 | 0.003 | 0.130 | 45.236 | LasR-defective |
| 19SJO | 0.040 | 0.021 | 0.004 | 0.014 | 0.035 | 0.019 | 0.015 | 0.000 | 0.043 | 0.036 | 0.031 | 0.029 | 2699.065 | Functional LasR |
| 19SJV | | | | | 0.109 | 0.073 | 0.032 | 0.000 | 0.004 | 0.004 | 0.003 | 0.026 | | LasR-defective |
| 19R | | | | | 0.079 | 0.167 | 0.029 | 0.002 | 0.003 | 0.003 | 0.004 | 0.049 | | LasR-defective |
| 19SV | | | | | 0.067 | 0.045 | 0.018 | 0.000 | 0.020 | 0.004 | 0.031 | 0.017 | | Functional LasR |
| 32SP | 0.000 | 0.000 | 0.000 | 0.000 | 0.045 | 0.049 | 0.000 | 0.000 | 0.000 | 0.000 | 0.000 | 0.001 | | LasR-defective, HAQ-neg |
| 32SB | 0.000 | 0.000 | 0.000 | 0.000 | 0.029 | 0.030 | 0.000 | 0.000 | 0.000 | 0.000 | 0.001 | 0.001 | 35.019 | LasR-defective, HAQ-neg |
| 32R | | | | | 0.110 | 0.240 | 0.040 | 0.010 | 0.003 | 0.005 | 0.004 | 0.059 | | LasR-defective |
| 34JS | | | | | 0.141 | 0.115 | 0.028 | 0.000 | 0.101 | 0.009 | 0.081 | 0.028 | | Functional LasR |
| 34JR | 0.001 | 0.001 | 0.000 | 0.001 | 0.080 | 0.229 | 0.000 | 0.043 | 0.000 | 0.035 | 0.001 | 0.176 | | LasR-defective |
| 57SJ | | | | | 0.061 | 0.098 | 0.015 | 0.000 | 0.029 | 0.005 | 0.056 | 0.016 | | Functional LasR |
| 57SP | | | | | 0.070 | 0.131 | 0.026 | 0.001 | 0.075 | 0.022 | 0.066 | 0.035 | | Functional LasR |
| 57RV | 0.000 | 0.000 | 0.000 | 0.000 | 0.092 | 0.240 | 0.000 | 0.042 | 0.000 | 0.015 | 0.001 | 0.099 | | LasR-defective |
| 57RP | | | | | 0.101 | 0.075 | 0.033 | 0.005 | 0.003 | 0.004 | 0.006 | 0.048 | | LasR-defective |
| 78RJ | | | | | 0.053 | 0.076 | 0.010 | 0.000 | 0.021 | 0.004 | 0.034 | 0.021 | | Functional LasR |
| 78RV | 0.001 | 0.001 | 0.000 | 0.005 | 0.027 | 0.044 | 0.132 | 0.116 | 0.012 | 0.032 | 0.021 | 0.150 | 321.465 | LasR-defective, RhlR-active |
| 78S | | | | | 0.068 | 0.116 | 0.006 | 0.000 | 0.021 | 0.006 | 0.038 | 0.021 | | Functional LasR |
| PG201 | 0.000 | 0.000 | 0.000 | 0.000 | 0.043 | 0.040 | 0.000 | 0.002 | 0.000 | 0.005 | 0.000 | 0.024 | | LasR-defective |
| PUPa3 | | | | | 0.716 | 0.023 | 0.008 | 0.000 | 0.055 | 0.005 | 0.078 | 0.012 | | Functional LasR |
| PA-CL501 | 0.015 | 0.003 | 0.001 | 0.005 | 0.083 | 0.040 | 0.000 | 0.000 | 0.000 | 0.000 | 0.000 | 0.000 | | Functional LasR, HAQ-negative |
| PA-CL502 | 0.019 | 0.032 | 0.001 | 0.005 | 0.173 | 0.020 | 0.000 | 0.000 | 0.000 | 0.000 | 0.000 | 0.000 | | Functional LasR, HAQ-negative |
| PA-CL504 | 0.019 | 0.004 | 0.001 | 0.005 | 0.026 | 0.029 | 0.000 | 0.000 | 0.000 | 0.000 | 0.000 | 0.001 | 21.637 | Functional LasR, HAQ-negative |
| PA-CL505 | 0.023 | 0.028 | 0.002 | 0.006 | 0.069 | 0.058 | 0.000 | 0.000 | 0.000 | 0.000 | 0.000 | 0.000 | | Functional LasR, HAQ-negative |
| PA-CL506b | 0.015 | 0.003 | 0.001 | 0.003 | 0.042 | 0.040 | 0.000 | 0.000 | 0.000 | 0.000 | 0.000 | 0.000 | | Functional LasR, HAQ-negative |
| PA-CL507 | 0.020 | 0.006 | 0.001 | 0.008 | 0.029 | 0.025 | 0.000 | 0.000 | 0.000 | 0.000 | 0.000 | 0.000 | 848.931 | Functional LasR, HAQ-negative |

| | | | | | | | | | | | | | | |
|---|---|---|---|---|---|---|---|---|---|---|---|---|---|---|
| PA-CL508 | 0.021 | 0.019 | 0.001 | 0.004 | 0.160 | 0.088 | 0.000 | 0.000 | 0.000 | 0.000 | 0.000 | 0.000 | | Functional LasR, HAQ-negative |
| PA-CL509 | | | | | 0.045 | 0.187 | 0.059 | 0.014 | 0.006 | 0.009 | 0.002 | 0.042 | | LasR-defective |
| PA-CL511 | | | | | 0.030 | 0.147 | 0.055 | 0.147 | 0.005 | 0.039 | 0.002 | 0.152 | | LasR-defective |
| PA-CL512 | 0.000 | 0.000 | 0.000 | 0.001 | 0.022 | 0.038 | 0.172 | 0.157 | 0.013 | 0.037 | 0.006 | 0.124 | 165.889 | LasR-defective, RhlR-active |
| PA-CL513 | 0.028 | 0.022 | 0.002 | 0.016 | 0.041 | 0.023 | 0.028 | 0.000 | 0.036 | 0.044 | 0.018 | 0.058 | 8017.829 | Functional LasR |
| PA-CL514 | | | | | 0.156 | 0.029 | 0.005 | 0.000 | 0.092 | 0.035 | 0.038 | 0.019 | | Functional LasR |
| PA-CL515 | | | | | 0.108 | 0.044 | 0.007 | 0.000 | 0.093 | 0.035 | 0.044 | 0.015 | | Functional LasR |
| PA-CL516 | | | | | 0.078 | 0.041 | 0.005 | 0.000 | 0.081 | 0.020 | 0.041 | 0.010 | | Functional LasR |
| PA-CL517 | | | | | 0.075 | 0.049 | 0.009 | 0.000 | 0.049 | 0.012 | 0.034 | 0.010 | | Functional LasR |
| PA-CL518 | | | | | 0.126 | 0.059 | 0.009 | 0.000 | 0.042 | 0.010 | 0.033 | 0.009 | | Functional LasR |
| PA-CL519 | | | | | 0.145 | 0.062 | 0.014 | 0.000 | 0.097 | 0.009 | 0.060 | 0.009 | | Functional LasR |
| PA-CL520 | | | | | 0.237 | 0.034 | 0.007 | 0.000 | 0.065 | 0.012 | 0.035 | 0.010 | | Functional LasR |
| PA-CL521a | 0.002 | 0.000 | 0.000 | 0.002 | 0.063 | 0.120 | 0.082 | 0.068 | 0.042 | 0.072 | 0.014 | 0.204 | 1608.202 | LasR-defective, RhlR-active |
| PA-CL521b | 0.000 | 0.000 | 0.000 | 0.012 | 0.078 | 0.108 | 0.080 | 0.057 | 0.044 | 0.074 | 0.012 | 0.203 | 1571.788 | LasR-defective, RhlR-active |
| PA-CL521c | 0.000 | 0.000 | 0.000 | 0.000 | 0.020 | 0.046 | 0.012 | 0.034 | 0.006 | 0.023 | 0.000 | 0.022 | 2720.812 | LasR-defective, RhlR-active |
| PA-CL522b | | | | | 0.531 | 0.208 | 0.089 | 0.001 | 0.035 | 0.016 | 0.035 | 0.081 | | LasR-defective |
| PA-CL523b | | | | | 0.702 | 0.129 | 0.052 | 0.000 | 0.022 | 0.010 | 0.018 | 0.052 | | LasR-defective |
| PA-CL524 | | | | | 0.092 | 0.115 | 0.050 | 0.000 | 0.014 | 0.017 | 0.024 | 0.051 | | LasR-defective |
| PA-CL527 | 0.018 | 0.006 | 0.001 | 0.003 | 13.605 | 1.253 | 0.000 | 0.000 | 0.000 | 0.000 | 0.000 | 0.001 | | Functional LasR, HAQ-negative |
| PA-CL528 | 0.037 | 0.010 | 0.003 | 0.004 | 0.224 | 0.133 | 0.000 | 0.000 | 0.000 | 0.000 | 0.001 | 0.000 | | Functional LasR, HAQ-negative |
| PA-CL529 | 0.022 | 0.008 | 0.002 | 0.003 | 0.282 | 0.036 | 0.000 | 0.000 | 0.000 | 0.000 | 0.000 | 0.001 | | Functional LasR, HAQ-negative |
| PA-CL532 | 0.035 | 0.015 | 0.004 | 0.021 | 0.043 | 0.040 | 0.022 | 0.001 | 0.082 | 0.115 | 0.032 | 0.076 | 1107.768 | Functional LasR |
| PA-CL534a | 0.019 | 0.014 | 0.001 | 0.005 | 0.035 | 0.014 | 0.000 | 0.000 | 0.000 | 0.000 | 0.000 | 0.000 | | Functional LasR, HAQ-negative |
| PA-CL534b | 0.017 | 0.009 | 0.002 | 0.003 | 0.046 | 0.039 | 0.000 | 0.000 | 0.000 | 0.000 | 0.001 | 0.000 | | Functional LasR, HAQ-negative |
| PA-CL534x | 0.021 | 0.008 | 0.002 | 0.003 | 0.044 | 0.057 | 0.000 | 0.000 | 0.000 | 0.000 | 0.000 | 0.000 | | Functional LasR, HAQ-negative |
| PA-CL535 | 0.017 | 0.014 | 0.002 | 0.004 | 0.096 | 0.008 | 0.000 | 0.000 | 0.000 | 0.000 | 0.000 | 0.000 | | Functional LasR, HAQ-negative |
| PA-CL542a | 0.022 | 0.023 | 0.002 | 0.003 | 0.101 | 0.022 | 0.000 | 0.000 | 0.000 | 0.000 | 0.001 | 0.000 | | Functional LasR, HAQ-negative |

| | | | | | | | | | | | | | |
|---|---|---|---|---|---|---|---|---|---|---|---|---|---|
| PA-CL542b | 0.003 | 0.003 | 0.001 | 0.001 | 0.026 | 0.050 | 0.000 | 0.000 | 0.000 | 0.000 | 0.001 | 0.000 | | Functional LasR, HAQ-negative |
| PA-CL545a | 0.019 | 0.007 | 0.002 | 0.004 | 0.145 | 0.144 | 0.000 | 0.000 | 0.000 | 0.000 | 0.000 | 0.000 | | Functional LasR, HAQ-negative |
| PA-CL547b | 0.019 | 0.007 | 0.001 | 0.003 | 0.064 | 0.053 | 0.000 | 0.000 | 0.000 | 0.000 | 0.000 | 0.000 | | Functional LasR, HAQ-negative |
| PA-CL549 | 0.016 | 0.013 | 0.002 | 0.005 | 0.069 | 0.094 | 0.000 | 0.000 | 0.000 | 0.000 | 0.000 | 0.000 | | Functional LasR, HAQ-negative |
| DCB1 | 0.000 | 0.000 | 0.000 | 0.000 | 0.132 | 0.113 | 0.000 | 0.000 | 0.000 | 0.003 | 0.000 | 0.005 | | LasR-defective, HAQ-negative |
| DCB2 | 0.000 | 0.000 | 0.000 | 0.000 | 0.070 | 0.057 | 0.000 | 0.000 | 0.000 | 0.005 | 0.000 | 0.013 | | LasR-defective |
| DCB3 | 0.000 | 0.000 | 0.000 | 0.000 | 0.027 | 0.008 | 0.055 | 0.129 | 0.016 | 0.176 | 0.000 | 0.000 | 30.029 | LasR-defective |
| DCB4 | 0.000 | 0.000 | 0.000 | 0.001 | 0.160 | 0.064 | 0.000 | 0.001 | 0.000 | 0.004 | 0.000 | 0.010 | | LasR-defective |
| DCB5 | | | | | 0.045 | 0.071 | 0.014 | 0.087 | 0.002 | 0.036 | 0.001 | 0.020 | | LasR-defective |
| DCB6 | | | | | 0.061 | 0.051 | 0.005 | 0.123 | 0.001 | 0.042 | 0.001 | 0.018 | | LasR-defective |
| DCB7 | | | | | 0.019 | 0.097 | 0.030 | 0.401 | 0.007 | 0.056 | 0.002 | 0.039 | | LasR-defective |
| DCB8 | | | | | 0.034 | 0.091 | 0.032 | 0.468 | 0.009 | 0.082 | 0.000 | 0.035 | | LasR-defective |
| DCB9 | 0.000 | 0.003 | 0.000 | 0.002 | 0.070 | 0.627 | 0.015 | 0.000 | 0.003 | 0.005 | 0.003 | 0.021 | | LasR-defective, HAQ-negative |
| DCB10 | 0.000 | 0.000 | 0.002 | 0.001 | 0.106 | 0.061 | 0.000 | 0.000 | 0.000 | 0.000 | 0.000 | 0.001 | | LasR-defective, HAQ-negative |
| DCB11 | 0.000 | 0.000 | 0.000 | 0.000 | 0.045 | 0.033 | 0.000 | 0.000 | 0.000 | 0.000 | 0.000 | 0.001 | | LasR-defective, HAQ-negative |
| DCB12 | 0.020 | 0.015 | 0.002 | 0.022 | 0.069 | 0.089 | 0.077 | 0.024 | 0.046 | 0.068 | 0.019 | 0.228 | 88.548 | Functional LasR |
| DCB13 | 0.052 | 0.014 | 0.007 | 0.018 | 0.047 | 0.046 | 0.013 | 0.019 | 0.063 | 0.176 | 0.016 | 0.119 | 23198.407 | Functional LasR, RhlR-active |
| DCB14 | | | | | 0.024 | 0.056 | 0.046 | 0.003 | 0.034 | 0.016 | 0.031 | 0.064 | | Functional LasR |
| DCB15 | 0.000 | 0.003 | 0.000 | 0.001 | 0.287 | 0.512 | 0.005 | 0.000 | 0.002 | 0.002 | 0.001 | 0.010 | | LasR-defective, HAQ-negative |
| DCB16 | | | | | 0.111 | 0.499 | 0.030 | 0.203 | 0.007 | 0.053 | 0.001 | 0.025 | | LasR-defective |
| DCB17 | | | | | 0.022 | 0.027 | 0.029 | 0.062 | 0.005 | 0.090 | 0.002 | 0.000 | | LasR-defective |
| DCB18 | 0.000 | 0.000 | 0.000 | 0.000 | 0.065 | 0.066 | 0.000 | 0.000 | 0.000 | 0.002 | 0.000 | 0.003 | | LasR-defective, HAQ-negative |
| DCB19 | 0.000 | 0.000 | 0.000 | 0.000 | 0.045 | 0.067 | 0.000 | 0.000 | 0.000 | 0.001 | 0.000 | 0.001 | | LasR-defective, HAQ-negative |
| DCB20 | 0.000 | 0.000 | 0.000 | 0.000 | 0.020 | 0.063 | 0.000 | 0.000 | 0.000 | 0.000 | 0.000 | 0.000 | | LasR-defective, HAQ-negative |
| DCB21 | | | | | 0.148 | 0.049 | 0.007 | 0.000 | 0.061 | 0.010 | 0.059 | 0.015 | | Functional LasR |
| DCB22 | 0.023 | 0.013 | 0.001 | 0.011 | 0.034 | 0.035 | 0.000 | 0.000 | 0.000 | 0.000 | 0.000 | 0.001 | 34.390 | Functional LasR, HAQ-negative |
| DCB23 | | | | | 2.019 | 0.037 | 0.010 | 0.000 | 0.046 | 0.008 | 0.056 | 0.013 | | Functional LasR |

| | | | | | | | | | | | | | | |
|---|---|---|---|---|---|---|---|---|---|---|---|---|---|---|
| DCB24 | | | | | 0.040 | 0.062 | 0.006 | 0.000 | 0.004 | 0.003 | 0.004 | 0.008 | | LasR-defective |
| DCB25 | | | | | 0.081 | 0.080 | 0.016 | 0.000 | 0.008 | 0.003 | 0.008 | 0.013 | | LasR-defective |
| DCB26 | 0.000 | 0.000 | 0.000 | 0.000 | 0.025 | 0.059 | 0.000 | 0.000 | 0.000 | 0.000 | 0.000 | 0.000 | | LasR-defective, HAQ-negative |
| DCB27 | 0.030 | 0.017 | 0.001 | 0.005 | 0.096 | 0.038 | 0.000 | 0.000 | 0.000 | 0.000 | 0.001 | 0.000 | | Functional LasR, HAQ-negative |
| DCB28 | | | | | 0.340 | 0.206 | 0.073 | 0.003 | 0.010 | 0.003 | 0.043 | 0.032 | | LasR-defective |
| DCB29 | 0.000 | 0.000 | 0.000 | 0.000 | 0.046 | 0.089 | 0.000 | 0.000 | 0.000 | 0.001 | 0.002 | 0.001 | | LasR-defective, HAQ-negative |
| DCB30 | | | | | 0.072 | 0.040 | 0.009 | 0.000 | 0.024 | 0.005 | 0.022 | 0.009 | | Functional LasR |
| DCB31 | 0.041 | 0.014 | 0.004 | 0.021 | 0.018 | 0.027 | 0.021 | 0.000 | 0.053 | 0.085 | 0.032 | 0.074 | 33.477 | Functional LasR |
| DCB32 | | | | | 0.166 | 0.062 | 0.013 | 0.000 | 0.049 | 0.013 | 0.057 | 0.020 | | Functional LasR |
| DCB33 | | | | | 1.089 | 0.159 | 0.058 | 0.000 | 0.034 | 0.013 | 0.036 | 0.059 | | LasR-defective |
| DCB34 | | | | | 0.083 | 0.083 | 0.065 | 0.000 | 0.030 | 0.015 | 0.012 | 0.060 | | Functional LasR |
| DCB35 | 0.001 | 0.000 | 0.000 | 0.000 | 0.018 | 0.053 | 0.000 | 0.000 | 0.000 | 0.000 | 0.000 | 0.000 | | LasR-defective, HAQ-negative |
| DCB36 | 0.000 | 0.000 | 0.000 | 0.000 | 0.017 | 0.084 | 0.045 | 0.038 | 0.007 | 0.112 | 0.001 | 0.000 | 38.886 | LasR-defective |
| DCB37 | | | | | 0.046 | 0.037 | 0.004 | 0.000 | 0.011 | 0.016 | 0.010 | 0.021 | | Functional LasR |
| DCB38 | 0.000 | 0.000 | 0.000 | 0.002 | 0.017 | 0.028 | 0.110 | 0.072 | 0.005 | 0.014 | 0.014 | 0.113 | 214.685 | LasR-defective |
| DCB39 | | | | | 0.149 | 0.145 | 0.067 | 0.001 | 0.017 | 0.010 | 0.013 | 0.045 | | LasR-defective |
| DCB40 | | | | | 0.070 | 0.058 | 0.020 | 0.000 | 0.036 | 0.006 | 0.031 | 0.014 | | Functional LasR |
| DCB41 | 0.035 | 0.016 | 0.002 | 0.011 | 0.025 | 0.025 | 0.000 | 0.000 | 0.000 | 0.000 | 0.002 | 0.000 | | Functional LasR, HAQ-negative |
| DCB42 | | | | | 0.343 | 0.037 | 0.005 | 0.000 | 0.055 | 0.008 | 0.058 | 0.011 | | Functional LasR |
| DCB43 | 0.042 | 0.019 | 0.004 | 0.021 | 0.026 | 0.018 | 0.016 | 0.003 | 0.044 | 0.043 | 0.037 | 0.057 | 4139.912 | Functional LasR |
| DCB44 | | | | | 0.045 | 0.019 | 0.036 | 0.000 | 0.044 | 0.023 | 0.059 | 0.030 | | Functional LasR |
| DCB45 | 0.007 | 0.007 | 0.000 | 0.001 | 0.034 | 0.028 | 0.000 | 0.000 | 0.000 | 0.000 | 0.000 | 0.000 | | Functional LasR, HAQ-negative |
| DCB46 | | | | | 0.082 | 0.021 | 0.048 | 0.000 | 0.071 | 0.022 | 0.039 | 0.029 | | Functional LasR |
| DCB47 | | | | | 0.040 | 0.021 | 0.094 | 0.031 | 0.016 | 0.023 | 0.004 | 0.080 | | LasR-defective |
| DCB48 | | | | | 0.073 | 0.039 | 0.092 | 0.001 | 0.042 | 0.031 | 0.019 | 0.041 | | Functional LasR |
| DCB49 | | | | | 0.128 | 0.016 | 0.066 | 0.000 | 0.095 | 0.019 | 0.069 | 0.027 | | Functional LasR |
| DCB50 | 0.001 | 0.002 | 0.000 | 0.016 | 0.025 | 0.036 | 0.108 | 0.160 | 0.008 | 0.022 | 0.009 | 0.158 | 319.179 | LasR-defective, RhlR-active |
| DCB51 | | | | | 0.070 | 0.012 | 0.042 | 0.000 | 0.072 | 0.015 | 0.058 | 0.021 | | Functional LasR |
| DCB52 | | | | | 0.065 | 0.021 | 0.036 | 0.000 | 0.062 | 0.022 | 0.069 | 0.033 | | Functional LasR |
| DCB53 | | | | | 0.047 | 0.027 | 0.047 | 0.000 | 0.028 | 0.018 | 0.028 | 0.039 | | Functional LasR |
| DCB54 | 0.000 | 0.000 | 0.000 | 0.006 | 0.036 | 0.077 | 0.081 | 0.053 | 0.009 | 0.026 | 0.009 | 0.090 | 26.229 | LasR-defective |

| | | | | | | | | | | | | | | |
|---|---|---|---|---|---|---|---|---|---|---|---|---|---|---|
| DCB55 | | | | | 0.048 | 0.028 | 0.083 | 0.005 | 0.033 | 0.019 | 0.035 | 0.068 | | LasR-defective |
| DCB56 | | | | | 0.029 | 0.030 | 0.076 | 0.012 | 0.011 | 0.013 | 0.012 | 0.072 | | LasR-defective |
| DCB57 | | | | | 0.063 | 0.037 | 0.134 | 0.006 | 0.023 | 0.017 | 0.017 | 0.061 | | LasR-defective |
| DCB58 | | | | | 0.074 | 0.047 | 0.148 | 0.042 | 0.030 | 0.025 | 0.009 | 0.067 | | LasR-defective |
| DCB59 | | | | | 0.051 | 0.033 | 0.142 | 0.113 | 0.022 | 0.020 | 0.006 | 0.134 | | LasR-defective |
| DCB60 | | | | | 0.056 | 0.042 | 0.067 | 0.000 | 0.052 | 0.028 | 0.046 | 0.070 | | Functional LasR |
| DCB61 | | | | | 0.272 | 0.032 | 0.642 | 0.003 | 0.207 | 0.024 | 0.286 | 0.066 | | Functional LasR |
| DCB62 | 0.030 | 0.016 | 0.004 | 0.011 | 0.032 | 0.073 | 0.026 | 0.008 | 0.046 | 0.111 | 0.006 | 0.101 | 26.847 | LasR-defective |
| DCB63 | | | | | 0.047 | 0.042 | 0.089 | 0.009 | 0.025 | 0.021 | 0.018 | 0.056 | | LasR-defective |
| DCB64 | | | | | 0.045 | 0.041 | 0.089 | 0.003 | 0.010 | 0.010 | 0.006 | 0.100 | | LasR-defective |
| DCB65 | | | | | 0.032 | 0.017 | 0.029 | 0.000 | 0.018 | 0.028 | 0.028 | 0.058 | | Functional LasR |
| DCB66 | | | | | 0.036 | 0.018 | 0.075 | 0.004 | 0.015 | 0.015 | 0.004 | 0.056 | | LasR-defective |
| DCB67 | | | | | 0.030 | 0.031 | 0.049 | 0.001 | 0.018 | 0.017 | 0.011 | 0.042 | | Functional LasR |
| DCB68 | | | | | 0.032 | 0.045 | 0.081 | 0.019 | 0.017 | 0.027 | 0.006 | 0.091 | | LasR-defective |
| DCB69 | | | | | 0.041 | 0.035 | 0.147 | 0.007 | 0.028 | 0.025 | 0.007 | 0.060 | | LasR-defective |
| DCB70 | | | | | 0.027 | 0.024 | 0.058 | 0.001 | 0.014 | 0.024 | 0.034 | 0.051 | | Functional LasR |
| DCB71 | | | | | 0.039 | 0.024 | 0.024 | 0.000 | 0.034 | 0.015 | 0.035 | 0.028 | | Functional LasR |
| DCB72 | | | | | 0.041 | 0.022 | 0.030 | 0.000 | 0.035 | 0.022 | 0.049 | 0.040 | | Functional LasR |
| DCB73 | | | | | 0.067 | 0.044 | 0.102 | 0.014 | 0.026 | 0.014 | 0.008 | 0.061 | | LasR-defective |
| DCB74 | 0.000 | 0.014 | 0.000 | 0.001 | 0.012 | 0.009 | 0.036 | 0.232 | 0.005 | 0.139 | 0.001 | 0.014 | 52.632 | LasR-defective, RhlR-active |
| DCB75 | | | | | 0.047 | 0.028 | 0.072 | 0.010 | 0.005 | 0.003 | 0.035 | 0.056 | | Functional LasR |
| DCB76 | 0.000 | 0.000 | 0.000 | 0.000 | 0.038 | 0.015 | 0.000 | 0.000 | 0.000 | 0.001 | 0.001 | 0.001 | | LasR-defective |
| DCB77 | | | | | 0.034 | 0.025 | 0.046 | 0.000 | 0.026 | 0.014 | 0.042 | 0.047 | | Functional LasR |
| DCB78 | | | | | 0.031 | 0.013 | 0.072 | 0.090 | 0.012 | 0.033 | 0.006 | 0.030 | | LasR-defective |
| DCB79 | | | | | 0.040 | 0.023 | 0.090 | 0.009 | 0.023 | 0.018 | 0.029 | 0.071 | | Functional LasR |
| DCB80 | | | | | 0.032 | 0.026 | 0.044 | 0.001 | 0.029 | 0.025 | 0.030 | 0.053 | | Functional LasR |
| DCB81 | | | | | 0.032 | 0.024 | 0.006 | 0.000 | 0.048 | 0.028 | 0.018 | 0.015 | | Functional LasR |
| DCB82 | | | | | 0.057 | 0.042 | 0.073 | 0.000 | 0.042 | 0.017 | 0.030 | 0.042 | | Functional LasR |
| DCB83 | | | | | 0.047 | 0.035 | 0.074 | 0.000 | 0.035 | 0.018 | 0.029 | 0.058 | | Functional LasR |
| DCB84 | | | | | 0.049 | 0.017 | 0.017 | 0.000 | 0.057 | 0.015 | 0.037 | 0.021 | | Functional LasR |
| DCB85 | 0.000 | 0.000 | 0.000 | 0.000 | 0.056 | 0.081 | 0.002 | 0.098 | 0.001 | 0.110 | 0.001 | 0.006 | 45.695 | LasR-defective |
| DCB86 | 0.000 | 0.001 | 0.000 | 0.003 | 0.023 | 0.053 | 0.134 | 0.076 | 0.021 | 0.052 | 0.012 | 0.151 | 354.323 | LasR-defective, RhlR-active |
| DCB87 | | | | | 0.068 | 0.024 | 0.045 | 0.000 | 0.065 | 0.013 | 0.082 | 0.029 | | Functional LasR |

| | | | | | | | | | | | | | |
|---|---|---|---|---|---|---|---|---|---|---|---|---|---|
| DCB88 | 0.036 | 0.017 | 0.003 | 0.020 | 0.022 | 0.018 | 0.010 | 0.000 | 0.030 | 0.031 | 0.012 | 0.023 | 4959.208 | Functional LasR |
| DCB89 | | | | | 0.032 | 0.025 | 0.069 | 0.006 | 0.021 | 0.017 | 0.018 | 0.080 | | Functional LasR |
| DCB90 | | | | | 0.048 | 0.015 | 0.025 | 0.000 | 0.051 | 0.018 | 0.034 | 0.020 | | Functional LasR |
| DCB91 | | | | | 0.044 | 0.063 | 0.093 | 0.815 | 0.015 | 0.090 | 0.001 | 0.120 | | LasR-defective |
| DCB92 | | | | | 0.050 | 0.040 | 0.048 | 0.001 | 0.052 | 0.047 | 0.055 | 0.099 | | Functional LasR |
| DCB93 | | | | | 0.023 | 0.019 | 0.003 | 0.002 | 0.001 | 0.001 | 0.002 | 0.003 | | LasR-defective |
| DCB94 | | | | | 0.036 | 0.017 | 0.023 | 0.001 | 0.035 | 0.029 | 0.046 | 0.039 | | Functional LasR |
| DCB95 | | | | | 0.039 | 0.009 | 0.084 | 0.219 | 0.016 | 0.167 | 0.002 | 0.000 | | LasR-defective |
| DCB96 | 0.026 | 0.039 | 0.002 | 0.011 | 0.031 | 0.009 | 0.000 | 0.000 | 0.000 | 0.000 | 0.001 | 0.000 | | Functional LasR, HAQ-negative |
| DCB97 | | | | | 0.062 | 0.021 | 0.035 | 0.001 | 0.066 | 0.026 | 0.058 | 0.037 | | Functional LasR |
| DCB98 | | | | | 0.040 | 0.026 | 0.047 | 0.007 | 0.007 | 0.011 | 0.003 | 0.035 | | LasR-defective |
| DCB99 | | | | | 0.126 | 0.061 | 0.159 | 0.001 | 0.090 | 0.065 | 0.094 | 0.091 | | Functional LasR |
| DCB100 | | | | | 0.071 | 0.022 | 0.050 | 0.000 | 0.054 | 0.023 | 0.030 | 0.029 | | Functional LasR |
| DCB101 | | | | | 0.037 | 0.020 | 0.014 | 0.000 | 0.045 | 0.022 | 0.035 | 0.026 | | Functional LasR |
| DCB102 | | | | | 0.051 | 0.035 | 0.061 | 0.001 | 0.010 | 0.008 | 0.010 | 0.044 | | Functional LasR |
| DCB103 | 0.001 | 0.005 | 0.000 | 0.000 | 0.026 | 0.015 | 0.000 | 0.001 | 0.000 | 0.004 | 0.000 | 0.003 | | LasR-defective |
| DCB104 | | | | | 0.060 | 0.032 | 0.015 | 0.001 | 0.042 | 0.021 | 0.049 | 0.032 | | Functional LasR |
| DCB105 | | | | | 0.048 | 0.018 | 0.018 | 0.000 | 0.040 | 0.015 | 0.032 | 0.021 | | Functional LasR |
| DCB106 | 0.000 | 0.000 | 0.000 | 0.000 | 0.047 | 0.072 | 0.000 | 0.002 | 0.000 | 0.005 | 0.001 | 0.004 | | LasR-defective |
| DCB107 | 0.020 | 0.013 | 0.000 | 0.001 | 0.029 | 0.023 | 0.000 | 0.000 | 0.000 | 0.000 | 0.001 | 0.000 | | Functional LasR, HAQ-negative |
| DCB108 | 0.000 | 0.000 | 0.000 | 0.006 | 0.038 | 0.049 | 0.089 | 0.029 | 0.015 | 0.023 | 0.014 | 0.101 | 267.161 | LasR-defective, RhlR-active |
| DCB109 | 0.000 | 0.001 | 0.000 | 0.007 | 0.038 | 0.064 | 0.119 | 0.086 | 0.025 | 0.053 | 0.018 | 0.153 | 295.151 | LasR-defective, RhlR-active |
| DCB110 | | | | | 0.027 | 0.027 | 0.063 | 0.017 | 0.015 | 0.029 | 0.005 | 0.079 | | LasR-defective |
| DCB111 | | | | | 0.057 | 0.036 | 0.113 | 0.011 | 0.022 | 0.015 | 0.012 | 0.105 | | LasR-defective |
| DCB112 | | | | | 0.071 | 0.036 | 0.115 | 0.005 | 0.037 | 0.023 | 0.027 | 0.084 | | Functional LasR |
| DCB113 | | | | | 0.044 | 0.042 | 0.096 | 0.034 | 0.015 | 0.026 | 0.006 | 0.108 | | LasR-defective |
| DCB114 | 0.026 | 0.014 | 0.001 | 0.005 | 0.033 | 0.026 | 0.000 | 0.020 | 0.000 | 0.003 | 0.000 | 0.002 | | LasR-defective |
| DCB115 | | | | | 0.029 | 0.010 | 0.063 | 0.009 | 0.012 | 0.019 | 0.001 | 0.018 | | LasR-defective |
| DCB116 | | | | | 0.062 | 0.042 | 0.139 | 0.000 | 0.023 | 0.000 | 0.015 | 0.000 | | Functional LasR |
| DCB117 | 0.000 | 0.000 | 0.000 | 0.000 | 0.022 | 0.106 | 0.067 | 0.015 | 0.019 | 0.093 | 0.001 | 0.000 | 41.823 | LasR-defective |
| DCB118 | 0.040 | 0.031 | 0.001 | 0.009 | 0.047 | 0.107 | 0.000 | 0.131 | 0.000 | 0.027 | 0.001 | 0.004 | | LasR-defective |
| DCB119 | | | | | 0.036 | 0.032 | 0.098 | 0.014 | 0.022 | 0.023 | 0.007 | 0.027 | | LasR-defective |

| | | | | | | | | | | | | | | |
|---|---|---|---|---|---|---|---|---|---|---|---|---|---|---|
| DCB120 | | | | | 0.030 | 0.022 | 0.000 | 0.002 | 0.000 | 0.001 | 0.001 | 0.002 | | LasR-defective |
| E90 | 0.000 | 0.000 | 0.000 | 0.002 | 0.018 | 0.021 | 0.020 | 0.015 | 0.020 | 0.015 | 0.039 | 0.029 | 707.692 | LasR-defective, RhlR-active |
| PA14 | 0.039 | 0.013 | 0.003 | 0.009 | 0.022 | 0.016 | 0.008 | 0.000 | 0.039 | 0.028 | 0.023 | 0.022 | 153.197 | Functional LasR |
| lasR PA14 | 0.001 | 0.000 | 0.000 | 0.002 | 0.019 | 0.026 | 0.042 | 0.019 | 0.005 | 0.018 | 0.003 | 0.044 | 40.558 | LasR-defective |

### Table S4. Whole genome sequence analyses

| # Assembly | Strain | Level | WGS | Chrs | BioSample | Largest contig | Total length | GC (%) | N50 | Origin | City | Country | Description | Source |
|---|---|---|---|---|---|---|---|---|---|---|---|---|---|---|
| GCA_004373595.1 | 18G | Scaffold | RWQV00000000 | undefined | SAMN10478256 | 1275041 | 6533293 | 66.3 | 960910 | Thouin's sandpit | Montreal | Canada | Contaminated soil (oil residue) | Déziel et al. 1996 |
| GCA_004372905.1 | 19R | Scaffold | RWQT00000000 | undefined | SAMN10478259 | 1304876 | 6536632 | 66.31 | 960677 | Thouin's sandpit | Montreal | Canada | Contaminated soil (oil residue) | Déziel et al. 1996 |
| GCA_004378685.1 | 19SJO | Contig | RWVK00000000 | undefined | SAMN10478257 | 1118602 | 6538449 | 66.3 | 745098 | Thouin's sandpit | Montreal | Canada | Contaminated soil (oil residue) | Déziel et al. 1996 |
| GCA_004373585.1 | 19SJV | Scaffold | RWQU00000000 | undefined | SAMN10478258 | 1154946 | 6536668 | 66.3 | 768130 | Thouin's sandpit | Montreal | Canada | Contaminated soil (oil residue) | Déziel et al. 1996 |
| GCA_004372645.1 | 19SV | Contig | RWQS00000000 | undefined | SAMN10478260 | 1118791 | 6540710 | 66.31 | 607825 | Thouin's sandpit | Montreal | Canada | Contaminated soil (oil residue) | Déziel et al. 1996 |
| GCA_004372875.1 | 32SB | Contig | RWQQ00000000 | undefined | SAMN10478262 | 1118249 | 6537026 | 66.3 | 615787 | Thouin's sandpit | Montreal | Canada | Contaminated soil (oil residue) | Déziel et al. 1996 |
| GCA_004372915.1 | 32SP | Contig | RWQR00000000 | undefined | SAMN10478261 | 1213340 | 6537249 | 66.3 | 486769 | Thouin's sandpit | Montreal | Canada | Contaminated soil (oil residue) | Déziel et al. 1996 |
| GCA_004372635.1 | 34JS | Contig | RWQP00000000 | undefined | SAMN10478263 | 1116732 | 6532148 | 66.3 | 302525 | Thouin's sandpit | Montreal | Canada | Contaminated soil (oil residue) | Déziel et al. 1996 |
| GCA_004372615.1 | 57RV | Contig | RWQN00000000 | undefined | SAMN10478265 | 1119279 | 6535060 | 66.3 | 943840 | Thouin's sandpit | Montreal | Canada | Contaminated soil (oil residue) | Déziel et al. 1996 |
| GCA_004372845.1 | 57SJ | Scaffold | RWQO00000000 | undefined | SAMN10478264 | 1693787 | 6460918 | 66.39 | 679751 | Thouin's sandpit | Montreal | Canada | Contaminated soil (oil residue) | Déziel et al. 1996 |
| GCA_004372595.1 | 78RV | Contig | RWQM00000000 | undefined | SAMN10478266 | 1119364 | 6534390 | 66.3 | 768454 | Thouin's sandpit | Montreal | Canada | Contaminated soil (oil residue) | Déziel et al. 1996 |
| GCA_004372585.1 | PA-CL501 | Contig | RWQL00000000 | undefined | SAMN10478267 | 1559844 | 6760284 | 66.27 | 675448 | Hospital sink | Montreal | Canada | Drain | Lalancette et al. 2017 |
| GCA_004372535.1 | PA-CL502 | Contig | RWQK00000000 | undefined | SAMN10478268 | 1456967 | 6760762 | 66.26 | 510299 | Hospital sink | Montreal | Canada | Drain | Lalancette et al. 2017 |
| GCA_004372525.1 | PA-CL504 | Contig | RWQJ00000000 | undefined | SAMN10478269 | 1554067 | 6756935 | 66.27 | 480257 | Hospital sink | Montreal | Canada | Drain | Lalancette et al. 2017 |
| GCA_004372505.1 | PA-CL505 | Contig | RWQI00000000 | undefined | SAMN10478270 | 1723252 | 6759636 | 66.27 | 650085 | Hospital sink | Montreal | Canada | Drain | Lalancette et al. 2017 |
| GCA_004372485.1 | PA-CL506b | Contig | RWQH00000000 | undefined | SAMN10478271 | 1442974 | 6763802 | 66.26 | 525188 | Hospital sink | Montreal | Canada | Drain | Lalancette et al. 2017 |
| GCA_004372495.1 | PA-CL507 | Contig | RWQG00000000 | undefined | SAMN10478272 | 1652205 | 6761925 | 66.24 | 510292 | Hospital sink | Montreal | Canada | Drain | Lalancette et al. 2017 |
| GCA_004372445.1 | PA-CL508 | Scaffold | RWQF00000000 | undefined | SAMN10478273 | 1733750 | 6763457 | 66.26 | 771247 | Hospital sink | Montreal | Canada | Drain | Lalancette et al. 2017 |
| GCA_004372825.1 | PA-CL509 | Contig | RWQE00000000 | undefined | SAMN10478274 | 1006921 | 6958392 | 66.01 | 630591 | Hospital sink | Montreal | Canada | Drain | Lalancette et al. 2017 |
| GCA_004372415.1 | PA-CL511 | Contig | RWQC00000000 | undefined | SAMN10478276 | 1091951 | 6961613 | 66.01 | 737267 | Hospital sink | Montreal | Canada | Drain | Lalancette et al. 2017 |
| GCA_004372395.1 | PA-CL512 | Contig | RWQB00000000 | undefined | SAMN10478277 | 1732063 | 6958739 | 66.01 | 681947 | Hospital sink | Montreal | Canada | Drain | Lalancette et al. 2017 |
| GCA_004372815.1 | PA-CL513 | Scaffold | RWQA00000000 | undefined | SAMN10478278 | 1446270 | 6716548 | 66.13 | 938712 | Hospital sink | Montreal | Canada | Drain | Lalancette et al. 2017 |
| GCA_004372375.1 | PA-CL514 | Contig | RWPZ00000000 | undefined | SAMN10478279 | 905139 | 6795047 | 66.2 | 579437 | Hospital sink | Montreal | Canada | Drain | Lalancette et al. 2017 |
| GCA_004372805.1 | PA-CL515 | Contig | RWPY00000000 | undefined | SAMN10478280 | 941291 | 6791188 | 66.2 | 600730 | Hospital sink | Montreal | Canada | Drain | Lalancette et al. 2017 |
| GCA_004372775.1 | PA-CL516 | Contig | RWPX00000000 | undefined | SAMN10478281 | 854620 | 6786301 | 66.21 | 425321 | Hospital sink | Montreal | Canada | Drain | Lalancette et al. 2017 |
| GCA_004372745.1 | PA-CL517 | Contig | RWPW00000000 | undefined | SAMN10478282 | 831540 | 7033028 | 66.01 | 640402 | Hospital sink | Montreal | Canada | Drain | Lalancette et al. 2017 |
| GCA_004372725.1 | PA-CL518 | Contig | RWPV00000000 | undefined | SAMN10478283 | 1105694 | 7028181 | 66.01 | 592523 | Hospital sink | Montreal | Canada | Drain | Lalancette et al. 2017 |
| GCA_004372345.1 | PA-CL519 | Contig | RWPU00000000 | undefined | SAMN10478284 | 1314378 | 7024063 | 66.02 | 591526 | Hospital sink | Montreal | Canada | Drain | Lalancette et al. 2017 |
| GCA_004372325.1 | PA-CL520 | Scaffold | RWPT00000000 | undefined | SAMN10478285 | 1419607 | 7030507 | 66.01 | 664650 | Hospital sink | Montreal | Canada | Drain | Lalancette et al. 2017 |
| GCA_004372315.1 | PA-CL521a | Contig | RWPS00000000 | undefined | SAMN10478286 | 1189888 | 7027941 | 66.09 | 311445 | Hospital sink | Montreal | Canada | Drain | Lalancette et al. 2017 |
| GCA_004372295.1 | PA-CL521b | Contig | RWPR00000000 | undefined | SAMN10478287 | 1783282 | 7010067 | 66.1 | 416245 | Hospital sink | Montreal | Canada | Drain | Lalancette et al. 2017 |
| GCA_004372275.1 | PA-CL522b | Scaffold | RWPQ00000000 | undefined | SAMN10478288 | 1135008 | 7004625 | 66.11 | 579094 | Hospital sink | Montreal | Canada | Drain | Lalancette et al. 2017 |
| GCA_004372705.1 | PA-CL524 | Contig | RWPP00000000 | undefined | SAMN10478289 | 1201753 | 7006586 | 66.11 | 811664 | Hospital sink | Montreal | Canada | Splash | Lalancette et al. 2017 |
| GCA_004372675.1 | PA-CL527 | Contig | RWPO00000000 | undefined | SAMN10478290 | 962520 | 6820892 | 66.21 | 429283 | Hospital sink | Montreal | Canada | Drain | Lalancette et al. 2017 |
| GCA_004372695.1 | PA-CL528 | Contig | RWPN00000000 | undefined | SAMN10478291 | 1962507 | 6816131 | 66.2 | 429687 | Hospital sink | Montreal | Canada | Drain | Lalancette et al. 2017 |
| GCA_004372255.1 | PA-CL529 | Contig | RWPM00000000 | undefined | SAMN10478292 | 1110037 | 6814260 | 66.2 | 534989 | Hospital sink | Montreal | Canada | Drain | Lalancette et al. 2017 |

| | | | | | | | | | | | | | | |
|---|---|---|---|---|---|---|---|---|---|---|---|---|---|---|
| GCA_004372205.1 | PA-CL532 | Contig | RWPL00000000 | undefined | SAMN10478293 | 1274754 | 6988055 | 65.97 | 558087 | Hospital sink | Montreal | Canada | Splash | Lalancette et al. 2017 |
| GCA_004372235.1 | PA-CL534a | Contig | RWPK00000000 | undefined | SAMN10478294 | 1522351 | 6818733 | 66.2 | 449994 | Hospital sink | Montreal | Canada | Faucet | Lalancette et al. 2017 |
| GCA_004378765.1 | PA-CL534x | Contig | RWPJ00000000 | undefined | SAMN10478295 | 1254195 | 6818156 | 66.2 | 510742 | Hospital sink | Montreal | Canada | Faucet | Lalancette et al. 2017 |
| GCA_004371955.1 | PA-CL542a | Contig | RWPI00000000 | undefined | SAMN10478296 | 1594899 | 6758636 | 66.27 | 615501 | Hospital sink | Montreal | Canada | Splash | Lalancette et al. 2017 |
| GCA_004371965.1 | PA-CL542b | Contig | RWPH00000000 | undefined | SAMN10478297 | 895351 | 6745722 | 66.27 | 340940 | Hospital sink | Montreal | Canada | Splash | Lalancette et al. 2017 |
| GCA_004372195.1 | PA-CL547b | Contig | RWPF00000000 | undefined | SAMN10478299 | 772370 | 6816421 | 66.21 | 414372 | Hospital sink | Montreal | Canada | Faucet | Lalancette et al. 2017 |
| GCA_004372145.1 | PA-CL549 | Contig | RWPE00000000 | undefined | SAMN10478300 | 1441280 | 6811068 | 66.21 | 563210 | Hospital sink | Montreal | Canada | Splash | Lalancette et al. 2017 |
| GCA_003838485.1 | PUPa3 | Contig | NSVI00000000 | undefined | SAMN07424021 | 771329 | 6333068 | 66.48 | 396213 | Rice rhizosphere | ND | India | Soil | Kumar et al. 2005 |
| GCA_003837775.1 | Rsan-ver | Contig | NSVJ00000000 | undefined | SAMN07424020 | 1311496 | 6841959 | 66.25 | 394571 | | Zurich | Switzerland | Soil | Guerra-Santos et al. 198[illegible] |
